# TIGIT-NECTIN2/3 signaling preserves ignorant CD8⁺ T cells for favorable immune checkpoint outcomes in HBV-related hepatocellular carcinoma

**DOI:** 10.64898/2026.05.06.723140

**Authors:** Pei-Ching Chang, Zhi-Chen Yan, Yung-Chih Hong, Kai-Ting Yu, Tze-Yun Hu, Pao-Shu Wu, Chen-Ching Lin, Tai-Ming Ko, Jinn-Moon Yang, Muh-Hwa Yang, Chun-Ying Wu, Jiunn-Chang Lin, Yan-Hwa Wu Lee

## Abstract

**Background:** Immune checkpoint inhibitors (ICIs) have revolutionized cancer therapy by restoring anti-tumor immunity. However, persistent antigen exposure drives T cell exhaustion, limiting the effectiveness of ICIs. Ignorant T cells are antigen-specific T cells that maintain a naïve state by regaining stem-like properties, allowing them to remain fully responsive to subsequent immunization. Virus-related hepatocellular carcinoma (HCC) demonstrates superior responses to ICIs compared to non-viral HCC, prompting us to investigate whether immunologically ignorant T cells exist in HBV-associated HCC and represent a promising target for improving immunotherapy outcomes.

**Methods:** Single-cell RNA sequencing (scRNA-seq) was performed on tumor tissues from patients with HBV-associated HCC. For validation, immunostaining was conducted on the discovery cohort and an independent cohort of 16 non-B non-C HCC and 22 HBV HCC. The enrichment of TIGIT and NECTIN3 in the proposed ignorant T cell was further validated using the TCGA database.

**Results:** scRNA-seq identified distinct HBV-infected HCC populations and revealed NECTIN3 upregulation in HBV-enriched subsets. CellChat analysis uncovered a novel NECTIN3-TIGIT tumor-immune interaction in HBV-enriched subsets, which shifted toward TIGIT-NECTIN2 as viral transcription declines. Trajectory analysis revealed the emergence of ignorant CD8⁺ T cells following T cell exhaustion. TIGIT-NECTIN2/3 interactions deliver a weak exhaustion signal. This allows T cells to survive and regain naïve-like properties as ignorant cells. Integration of bulk RNA-seq data identified CD24, STMN1, and EZH2 as potential biomarkers of ignorant CD8⁺ T cells.

**Conclusions:** TIGIT-NECTIN2/3 interactions present a promising axis for preserving immunologically ignorant T cells and sustaining ICI responsiveness in HBV-associated HCC.

## Introduction

Chronic hepatitis B virus (HBV) infection accounts for ∼55%-80% of hepatocellular carcinoma (HCC) (1, 2). Although HBV vaccination programs have reduced new infections, chronic HBV continues to affect ∼350 million people worldwide (3). Traditional treatments have limited efficacy against unresectable advanced HCC (4). While immune checkpoint inhibitors (ICIs), such as atezolizumab/bevacizumab (anti-PD-L1/anti-VEGF) (5) or durvalumab/tremelimumab (anti-PD-L1/anti-CTLA-4) (6) has improved outcomes, immune escape through immune checkpoints LAG-3 (7), TIM3/HAVCR2 (8), and TIGIT (9) remains a challenge.

Persistent antigen exposure, as seen in cancer and chronic viral infections, drives T cell exhaustion, characterized by loss of effector functions, reduced proliferative capacity, transcriptional reprogramming, and upregulation of inhibitory receptors, including PD-1, CTLA-4, LAG-3, TIM3, and TIGIT (10, 11). Single-cell RNA sequencing (scRNA-seq) has identified subsets of exhausted T cells, including a progenitor subset (T_PEX_) that expresses TCF-1 and retains stem-like and proliferative properties, allowing them to self-renew and give rise to intermediate and terminally exhausted T cells (12), potentially explaining the favorable responses to ICIs observed in HBV-associated HCC (13). Interestingly, clinical observations have demonstrated favorable responses to ICIs in HBV-associated HCC (14, 15). These features position HBV-specific T cells in HBV-positive HCC tumors as a valuable model for investigating tumor-specific T cell responses.

Effective engagement of T cells by antigen-presenting cells is critical for initiating cytotoxic responses, whereas sustained T cell receptor (TCR) signaling leads to exhaustion. Recently, sequential stages of CD8⁺ T cell states under persistent antigen exposure in cancer and viral infections have been postulated, with the framework extended to include immunological ignorance as a novel stage (16). In this conceptual model, diseased cells fail to provide sufficient antigenic stimulation, allowing CD8⁺ T cells to re-acquire a naïve-like stemness phenotype referred to as ignorant T cells. These “ignorant” T cells resemble T_PEX_ cells and are distinct from tolerance, anergy, or terminal exhaustion (16), providing a potential mechanistic basis for the favorable ICI responses observed in virus-associated HCC (14). In HBV-associated HCC, CD8⁺ T cells encountering HBV-infected tumor cells may fail to activate fully and instead enter a state of immunological ignorance, while functionally suppressed, retain the capacity to regain effector function upon immune checkpoint blockade. Thus, a deeper understanding of immunosuppressive mechanisms, particularly the crosstalk between HBV-infected HCC cells and T cells, is critical for advancing immunotherapy strategies. The crosstalk between cells in HCC tumor microenvironment (TME) is highly complex, comprising both malignant hepatocytes and diverse infiltrating immune cells that together shape tumor progression and therapeutic response. Many scRNA-seq studies of HCC have provided important insights into tumor-infiltrating and circulating immune cells (17–21). Interaction analyses have largely emphasized immune-immune crosstalk, particularly between myeloid cells and T cells (21, 22). Only a limited number of studies have resolved HCC cells of HBV etiology or their potential impact on the immune microenvironment. For example, Ho *et al.* delineated HCC clusters and characterized the immunosuppressive landscape of HBV-associated HCC, but their interaction analysis demonstrated that macrophages suppress tumor-infiltrating T cells through TIGIT-NECTIN2 axis (22). Guo *et al.* revealed the heterogeneity of HCC and reported HCC-fibroblast crosstalk (23). More recently, Liu *et al.* deciphered HBV transcripts in HCC and compared the interactomes of HBV- and non-HBV-associated tumors, showing that myeloid cells primarily engaged T/NK cells through cytokine signaling, including the immunosuppressive interaction in HBV-HCC and the anti-tumor interaction in non-HBV-HCC (24). Despite these advances, the single-cell characteristics of HBV-infected HCC cell interactions remain largely unexplored, and their contribution to shaping immune responses, which may underlie the favorable outcomes of immunotherapy in virus-associated HCC (14), remains poorly understood. To address this, we performed scRNA-seq on clinical specimens from HBV-associated HCC patients, aiming to capture both tumor and immune compartments and to uncover tumor-immune interactions linked to the favorable immunotherapeutic responses in virus-associated HCC.

## Materials and Methods

### Human sample collection

Human hepatocellular carcinoma (HCC) specimens were collected at Mackay Memorial Hospital, Taiwan, under Institutional Review Board (IRB)-approved protocols (Approval No. 13MMHIS269), with written informed consent obtained from all patients before sample collection. In 2020, four fresh HCC specimens positive for hepatitis B virus (HBV) and negative for hepatitis C virus (HCV) and hepatitis D virus (HDV) were obtained. Each was divided for clinical and research use: one portion was formalin-fixed and paraffin-embedded (FFPE) for histopathological diagnosis, and the other processed for single-cell preparation. Additionally, between 2014 and 2023, 38 FFPE HCC specimens were collected, including 22 HBV-positive cases with serum HBV DNA levels > 200 IU/mL and 16 non-B non-C (NBNC) HCC cases.

### Single-cell data analysis

Pseudotime trajectory analysis was performed using Monocle 2 (25) with default parameters in Partek Flow. For HCC cells and tumor-infiltrating T cells, all expressed genes from each cell type were used to construct the pseudotime trajectory. Reversed Graph Embedding was applied to project cells onto a low-dimensional manifold, enabling the ordering of cells along a trajectory and the identification of branch points representing potential cell fate decisions. Cell-cell communication was analyzed by evaluating ligand-receptor interactions using the CellChat R package (v2.1.0) (26) with default parameters, with the trimean approach requiring 25% of the cells in a population to express a specific ligand or receptor to be considered for statistical testing.

### Immunohistochemical staining

FFPE sections were processed on the automated Ventana Benchmark XT (Ventana Medical Systems, Inc.) using a NECTIN3-specific antibody (Abcam) at room temperature. Staining intensity was evaluated using an H-score based on a four-tier scale: negative (0), weak (1), moderate (2), and strong (3).

### Multiplex immunofluorescence staining

Multiplex staining of FFPE tissue was performed using the Opal™ Opal™ 6-PLEX MANUAL DETECTION KIT (Akoya Biosciences), according to the manufacturer’s instructions. Detailed procedures are described in the Supplementary Methods.

## Results

### Single-cell analysis of HBV-associated hepatocellular carcinoma

Given the favorable clinical responses to ICIs observed in HBV-associated HCC (14, 15), understanding the unique features of TME helps restore anti-tumor immune activity. To date, single-cell studies of HCC TME have largely focused on delineating the immune landscape and immune-immune crosstalk (17, 27), leaving the interaction characteristics of HBV-infected HCC cells poorly defined. Elucidating how HBV-infected HCC cells interact with immune cells helps explain the superior responses of virus-related HCC to ICIs compared with non-virus-related HCC. Accordingly, we focus on characterizing HCC-T cell interactions, with particular emphasis on immune checkpoint pathways.

To this end, we performed scRNA-seq on four HBV-associated HCC specimens as a discovery cohort (Table S1). Longitudinal serum HBV DNA levels and sampling time points are presented in Supplementary Fig. 1. Following initial quality control, doublet removal, and batch-effect correction, we obtained 22,468 single-cell transcriptomes (Table S2). Unsupervised graph-based clustering revealed 13 distinct cell clusters (Fig. 1A). Cell types were initially annotated using CIPR R package and subsequently confirmed by cell type-specific markers (Supplementary Fig. 2), identifying B cells, T cells, natural killer (NK) cells, macrophages, endothelial cells, stellate cells, and HCC cells (Fig. 1A). Notably, cell clusters contained single cells from multiple cases admixed together (Fig. 1B), demonstrating a relatively low level of inter-tumoral heterogeneity. Analysis of 3,732 HCC cells revealed six distinct HCC clusters (Fig. 1C). We then mapped sequencing reads that did not align to the human reference genome (hg38) to the HBV genome. Among the four major HBV genes, polymerase (gp1), surface antigen (HBs-Ag; gp2), X protein (HBx; gp3), and core protein (HBc; gp4), only HBs-Ag and HBx transcripts were detected (Fig. 1D). Viral transcript mapping defined HBV-infected and uninfected cells and identified HCC-c1 and HCC-c2 as HBV^High^, HCC-c3, HCC-c4, and HCC-c5 as HBV^Inte^, and HCC-c6 as HBV^Low^ (Fig. 1E). To uncover the key immune checkpoint ligands associated with HBV infection, we examined the expression of immunosuppressive ligands in HCC clusters (Fig. 1F). Interestingly, among the examined ligands, only NECTIN3 expression showed a strong correlation with HBV-enriched clusters (HCC-c1/c2) (r=0.75; Fig. 1G, upper panel), whereas NECTIN2 (Fig. 1G, lower panel), as well as other ligands, did not display a significant association. This result indicates a potential involvement of the TIGIT-NECTIN3 axis in HBV-related HCC.

**Fig. 1.**
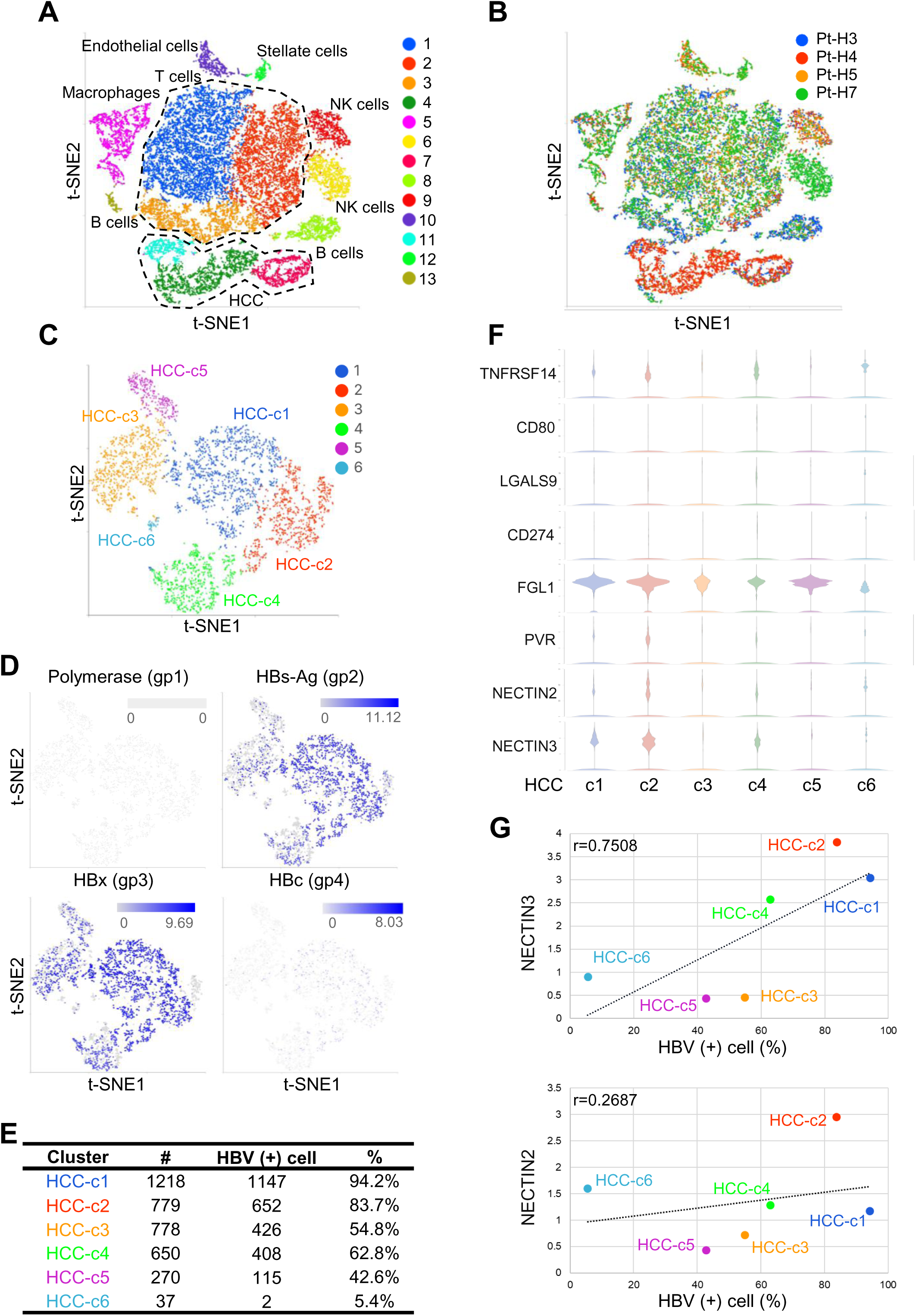
Single-cell transcriptomic profiling identifies HBV-infected hepatocellular carcinoma (HCC) cells in HBV-associated HCC specimens. (A) Cell-type identification of single cells from HCC tumors. (B) Cell clusters annotated by identity across four patients. (C) t-SNE projection of HCC cells from HBV-associated HCC tumors, revealing six distinct clusters. (D) Expression of HBV genes polymerase (gp1), surface antigen (HBs-Ag; gp2), X protein (HBx; gp3), and core protein (HBc; gp4) across clusters. (E) Proportions of HBV-positive HCC cells within each cluster. (F) Violin plots showing the expression of immunosuppressive ligands across HCC clusters. (G) Correlation of NECTIN2 and NECTIN3 expression with HBV-positive cell percentages.

### Single-cell landscape of T cell compartment in HBV-associated HCC

CD8⁺ T cells play a critical role in viral clearance but become functionally impaired during chronic infection, leading to immune exhaustion and reduced effectiveness of ICI therapy (28, 29). However, exhausted CD8⁺ T cells may retain functional plasticity and reacquire naïve- or stemness-like features, exemplified by progenitor exhausted (T_PEX_) cells, which likely serve as the cellular basis for superior response to ICIs (13, 30). To uncover novel progenitor populations, we first delineated T cell subtypes by performing unsupervised clustering analysis of cytotoxic lymphocytes, including T cells and NK cells. A total of 15,023 lymphocytes were grouped into eight distinct clusters (Fig. 2A). Using cell type-specific markers (Supplementary Fig. 3), we defined four CD8⁺ T cell subsets, three CD4⁺ T cell subsets, one of which represented regulatory T cells (Tregs), and one NK cell cluster (Fig. 2A). The data revealed inter-patient variation in immune cell composition, with CD4^+^ and CD8^+^ T cells being the most abundant populations (Fig. 2B). To validate the scRNA-seq results, we performed multiplex immunofluorescence staining on formalin-fixed paraffin-embedded (FFPE) tissues from the four scRNA-seq specimens (Fig. 2C) and observed a comparable proportion of CD4⁺ and CD8⁺ T cells within the TME (Fig. 2D). We further examined the expression of key immunosuppressive molecules within CD8⁺ T, CD4⁺ T, Treg, and NK cell clusters (Fig. 2E), and found that CTLA-4 and TIGIT were highly expressed in Treg, while TIGIT was also expressed in clusters 1 and 4 of CD8⁺ T cells, with greater enrichment in cluster 4 (Fig. 2E). Our data suggest that chronic HBV infection involves immunosuppressive pathways beyond canonical PD-1-dependent T cell exhaustion. The enrichment of TIGIT in T cells further supports the notion that immunosuppression of CD8⁺ T cells in HBV-associated HCC may be mediated via the TIGIT-NECTIN axis.

**Fig. 2.**
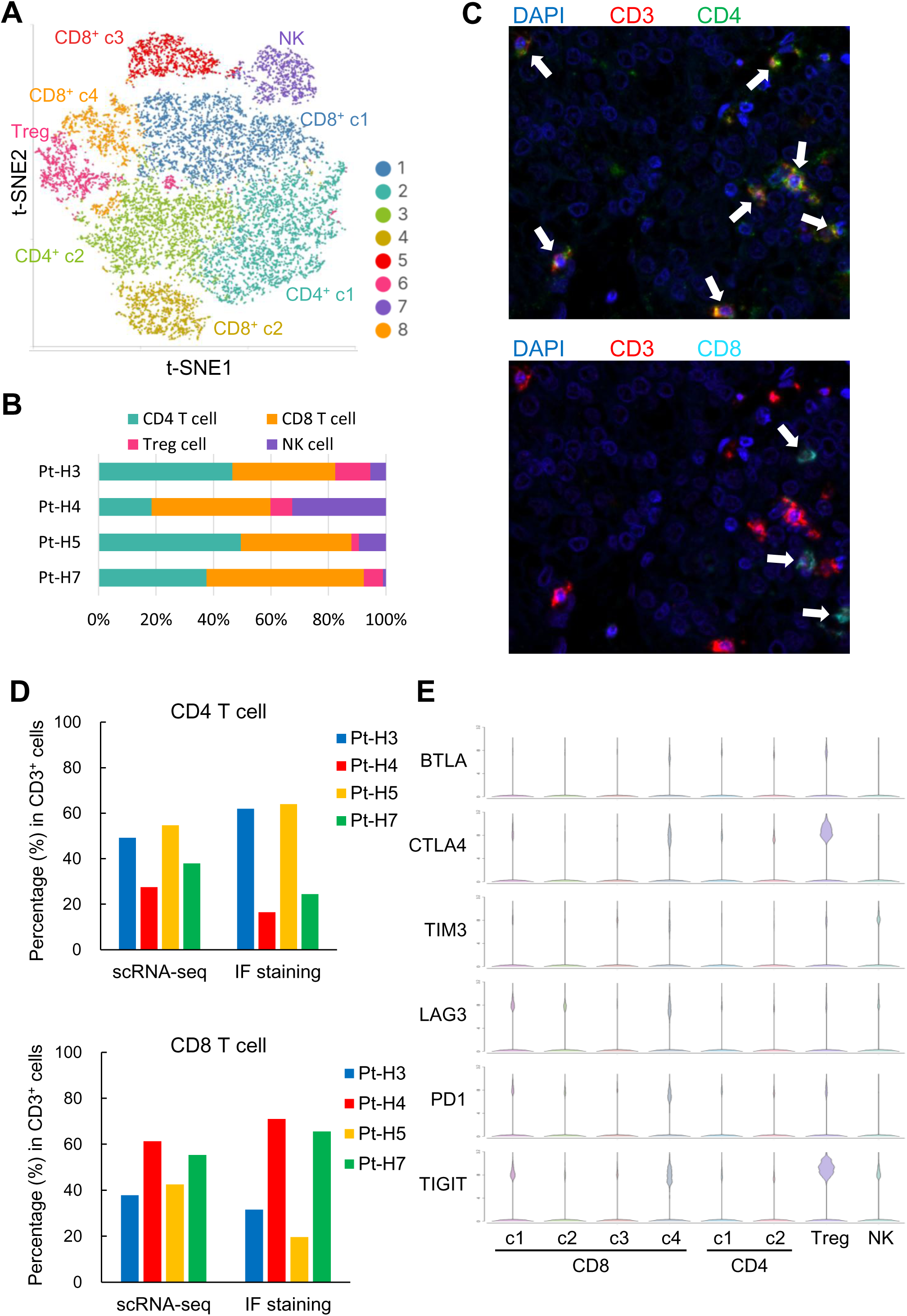
Dissection of tumor-infiltrating T cell clusters in HBV-associated HCC. (A) t-SNE projection of T cells from HBV-associated HCC tumors, revealing eight distinct clusters: CD4 (c1), CD4 (c2), Treg, CD8 (c1), CD8 (c2), CD8 (c3), CD8 (c4), and natural killer (NK) cells. (B) Proportion of T cell subsets and NK cells identified in each HBV-associated HCC tumor. (C) Representative images of multiplex immunofluorescence staining of anti-CD3, CD4, and CD8. (D) Quantification of CD4⁺ and CD8⁺ T cell populations among CD3⁺ T cells. (E) Violin plots showing the expression of immunosuppressive markers across T cell clusters.

### Mapping CD8⁺ T cell subsets reveal high TIGIT expression in progenitor exhausted T cell

To elucidate T cell subtypes with limited sample size, we utilized the ProjecTILs, which provides comprehensive human CD4⁺ (31) and CD8⁺ (32) tumor-infiltrating lymphocyte (TIL) atlases (Fig. 3A, upper panel), to integrate T cells from our scRNA-seq dataset onto defined T cell subsets. The CD8^+^ TIL atlas includes Naive-like T cells, central memory (T_CM_), effector memory (T_EM_), T_EMRA_ (effector memory re-expressing CD45RA), progenitor exhausted T cells (T_PEX_), terminally exhausted T cells (T_EX_), and mucosal-associated invariant T cells (T_MAIT_) (Fig. 3A, upper left panel) (32). The CD4^+^ TIL atlas comprises Naive-like T cells, T follicular helper (Tfh) cells, Th17 helper cells, T regulatory (Treg) cells, cytotoxic CD4^+^ T cells expressing EOMES and GZMK (CTL_EOMES), cytotoxic CD4^+^ T cells expressing GNLY (CTL_GNLY), and cytotoxic CD4^+^ T cells with an exhaustion phenotype (CTL_Exh) (Fig. 3A, upper right panel) (31). After projecting T cell subsets defined by ProjecTILs TIL atlases onto our CD4⁺ and CD8⁺ T cell clusters, we observed diverse proportions of CD8⁺ subsets, including naive-like T cells, T_CM_, T_EM_, T_EMRA_, T_PEX_, T_EX,_ and T_MAIT_ cells (Fig. 3A, lower left panel and Fig. 3B), whereas CD4⁺ T cells were predominantly naive-like T cells and Treg (Fig. 3A, lower right panel and Fig. 3C). Consistent with previous studies, our analysis confirmed the expression of immunosuppressive molecules in T_EX_ cells (33). Notably, we further identified high TIGIT expression, together with PD-1 and CTLA-4, in T_PEX_ cells in HBV-associated HCC (Fig. 3D).

**Fig. 3.**
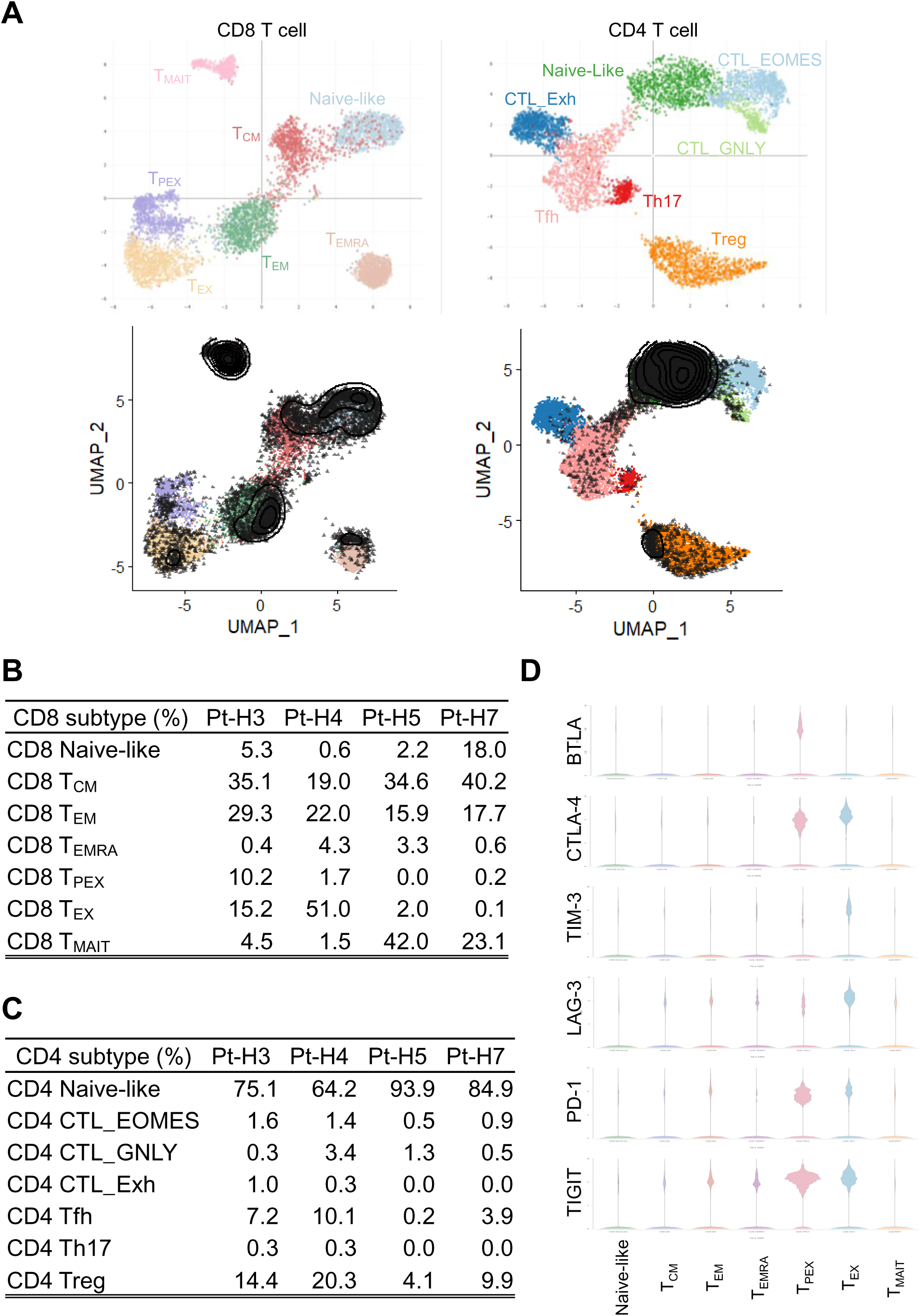
Inhibitory receptor expression in CD8^+^ T cell subsets in HBV-associated HCC. (A) t-SNE plots of T cells from scRNA-seq data. Upper panel: ProjecTILs-based annotation of CD8⁺ (left) and CD4⁺ (right) T cells. Lower panels: contour plots showing cell density distributions for the corresponding groups. CD8⁺ T cells were classified into naïve-like, central memory (T_CM_), effector memory (T_EM_), effector memory re-expressing CD45RA (T_EMRA_), progenitor exhausted (T_PEX_), terminally exhausted (T_EX_), and mucosal-associated invariant T cells (T_MAIT_). CD4⁺ T cells were primarily annotated as naïve-like and regulatory T cells (Treg). (B, C) Proportions of CD8⁺ (B) and CD4⁺ (C) T cell subsets. (D) Violin plots showing the expression of immunosuppressive markers BTLA, CTLA-4, TIM3, LAG-3, PD-1, and TIGIT across T cell subsets.

### Cell-state transition trajectory of tumor-infiltrating CD8⁺ T cell subpopulations

To delineate the evolution of CD8⁺ T cell within the TME of HBV-associated HCC, we performed pseudotime trajectory analysis of tumor-infiltrating CD8⁺ T cells and identified five distinct states (Fig. 4A). Projection of ProjecTILs-defined CD8⁺ T cell subsets onto these states revealed that state 1 was enriched for naive-like T cells and T_CM_ cells; state 2 was primarily composed of T_CM_ cells; state 3 included T_CM_ and T_EM_, T_EMRA_, T_PEX_, and T_EX_ cells, along with a minor population of T_MAIT_ cells; state 4 contained T_CM_ and T_EM_ cells, with a small proportion of T_MAIT_ cells; and state 5 included T_CM_, T_EM_, and was predominantly enriched in T_MAIT_ cells (Fig. 4B and Table S3). Integrating ProjecTILs-based annotation with trajectory analyses revealed a transition of CD8⁺ T cells from a naive-like state (state 1) toward two distinct trajectories: mucosal-associated specialization (T_MAIT_, state 5) and functional exhaustion (states 2 to 4), encompassing an admixture of T_EM_, T_EMRA_, T_PEX_, and T_EX_ (Fig. 4B, 4C). Notably, the ProjecTILs-defined T_PEX_ population emerged at the end of the exhaustion trajectory, rather than occupying an intermediate position between naive-like T cells and T_EX_. These T_PEX_-like cells expressed both T_PEX_ markers, including stem-like marker TCF-1 and exhaustion markers PD-1 and TOX, as described in chronic viral infection models (33), as well as naïve T cell markers, such as CCR7, CD62L, IL7R, CD27, and CD28) (34), identifying them as immunologically ignorant T cells, as proposed by Galluzzi *et al* (16). Since naïve-like ignorant CD8⁺ T cells may help explain the favorable responsiveness to ICIs observed in HBV-associated HCC compared to non-viral HCC (14), and the biomarker for ignorant CD8⁺ T cells remains unknown, we aimed to identify candidate biomarkers. To this end, we projected the CD8⁺ T cell clusters defined in Fig. 2A onto the pseudotime states. We found that CD8⁺ c4 was primarily located at the terminal end of state 3 (Fig. 4F), corresponding to the location of T_PEX_-like ignorant CD8⁺ T cells. Next, we performed differential gene expression analysis to define CD8⁺ c4 cluster-specific biomarkers. To identify the most highly expressed CD8⁺ c4 biomarkers, bulk RNA-seq was performed on tumor and adjacent non-tumor tissues from the four scRNA-seq HCC cases, as well as four recurrent samples (Supplementary Fig. 1), and cluster-specific biomarkers (tumor/adjacent ≥5-fold) were ranked by their expression levels (Table S4). The top 10 cluster-specific biomarkers were visualized using violin plots, highlighting CD24, STMN1, and EZH2 as potential biomarkers for the immunologically ignorant population (Fig. 4G).

**Fig. 4.**
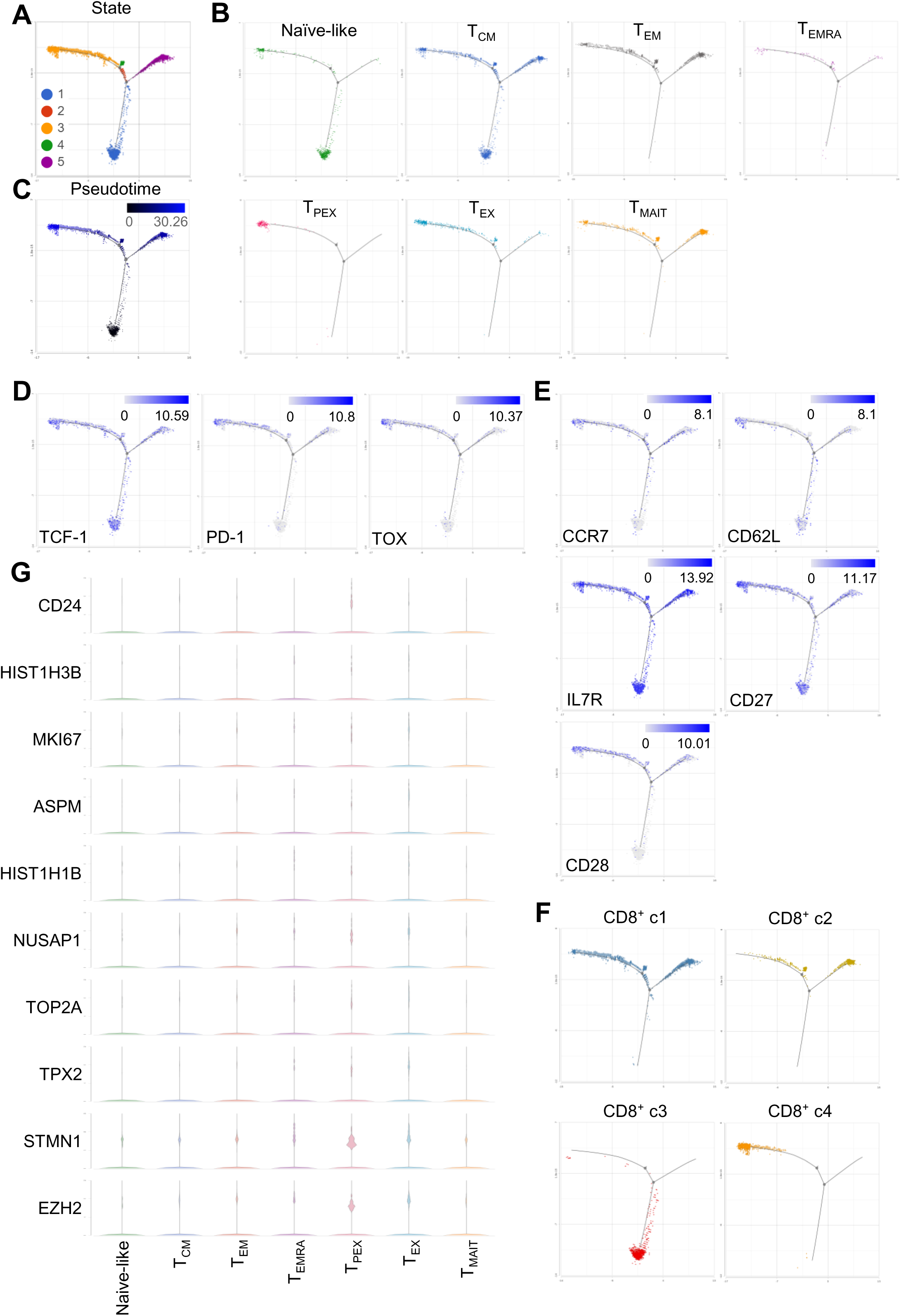
Pseudotime trajectory analysis of CD8⁺ T cells in HBV-associated HCC. (A) Monocle 2 trajectory of CD8⁺ T cells colored by state. (B) ProjecTILs-defined CD8⁺ T cell subtypes in each trajectory state. (C) Pseudotime trajectory analysis of CD8⁺ T cell. (D) Expression of T_PEX_ markers TCF-1, PD-1, and TOX along the trajectory. (E) Expression of Naïve T cell markers CCR7, CD62L, IL7R, CD27, and CD28 along the trajectory. (F) CD8⁺ T cell clusters in each trajectory state. (G) Violin plots showing the expression of top 10 CD8⁺ c4 cluster-specific biomarkers.

### Cell-cell communication between HBV-associated HCC cells and CD8⁺ T cells elucidate non-canonical immunosuppression in HBV-associated HCC

Crosstalk between HBV-infected HCC cells and immune cells may support the emergence of naïve-like ignorant CD8⁺ T cells. To investigate HBV-associated immunosuppressive mechanisms, we applied CellChat analysis (26) to examine cell-cell interactions between HCC clusters defined in our scRNA-seq dataset (Fig. 1C) and functionally distinct T cell subsets defined by ProjecTILs across patient specimens. Focusing on immunosuppressive ligand-receptor pairs, we identified TIGIT-NECTIN interactions as a dominant axis of suppression between HBV-enriched HCC clusters and CD8⁺ T cells, as well as Treg cells (Fig. 5A). In contrast, no significant interactions were observed between CD4⁺ T cells and HCC clusters, likely attributed to the predominance of naïve phenotypes within the CD4⁺ population (Fig. 3).

**Fig. 5.**
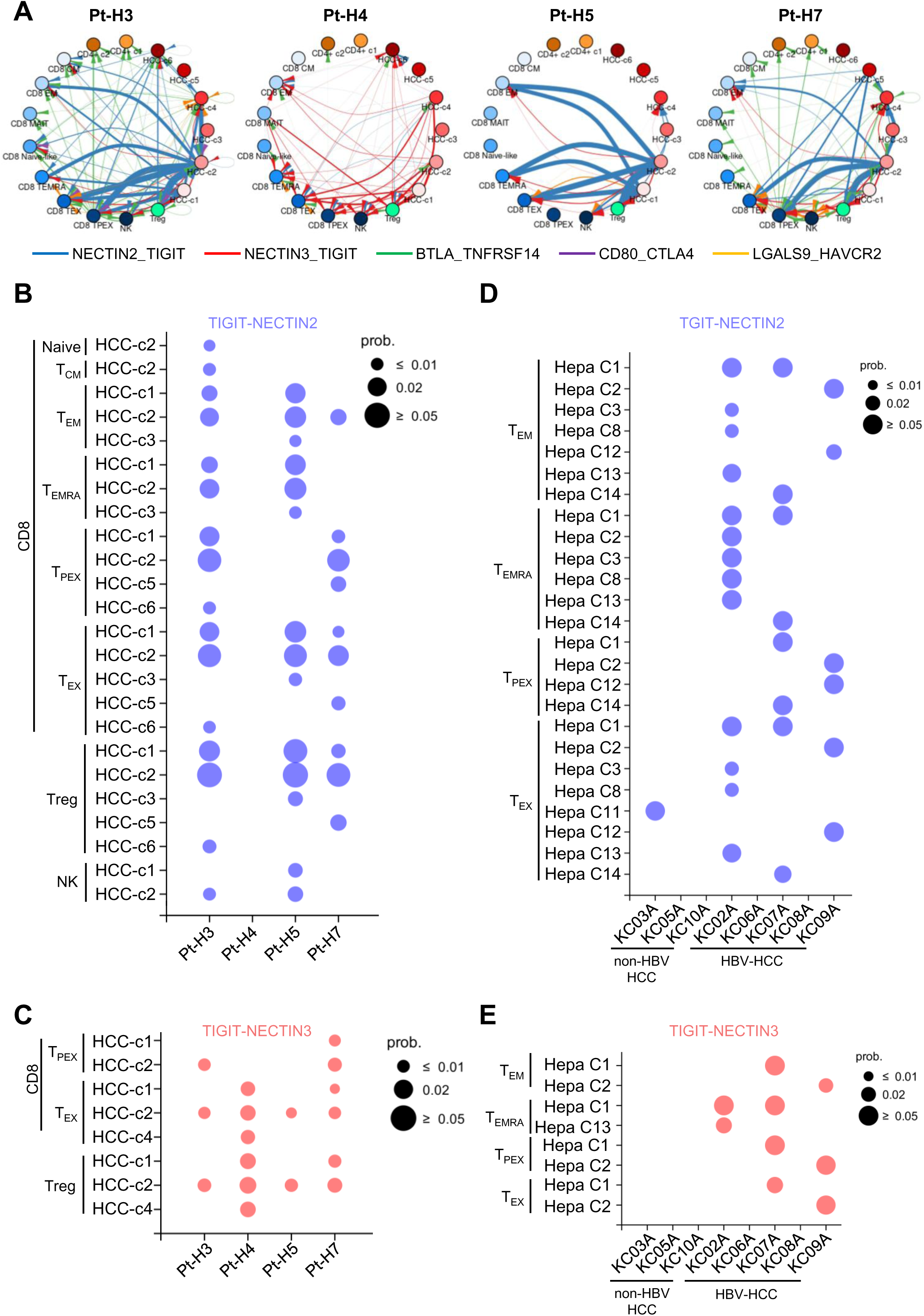
Deciphering TIGIT-NECTIN interactions between T cells and HCC cells in individual patient specimens. (A) Cell-cell communication networks involving key immunosuppressive molecules, including BTLA, CTLA-4, TIM3/HAVCR2, LAG3, PD-1, and TIGIT, and their corresponding ligands in HBV-associated HCC. (B, C) TIGIT-NECTIN2 (B) and TIGIT-NECTIN3 (C) interactions between ProjecTILs-defined T cell subtypes and HCC clusters. (D, E) TIGIT-NECTIN2 (D) and TIGIT-NECTIN3 (E) interactions between ProjecTILs-defined CD8^+^ T cell subtypes and HCC clusters in GSE282343 dataset.

Consistent with a previous report showing TIGIT-NECTIN2-mediated suppression by tumor-associated macrophages (TAMs) (22), we observed TIGIT-NECTIN2 interactions between T_EM_, T_EMRA_, T_PEX_, and T_EX_ CD8⁺ T cell and HCC-c1 and HCC-c2, and to a lesser extent with HCC-c3, HCC-c5, and HCC-c6 cells (Fig. 5B). Notably, TIGIT-NECTIN3 interaction was selectively detected between T_PEX_ CD8⁺ T cells and HCC-c1 and HCC-c2 clusters, as well as between T_EX_ CD8⁺ T cell and Treg cells with HCC-c1, HCC-c2, and HCC-c4 (Fig. 5C). Given that ProjecTILs-defined T_PEX_ cells represent an immunologically ignorant CD8⁺ T cell population in this study, the preferential engagement of NECTIN3-TIGIT signaling between HCC cells and ProjecTILs-defined T_PEX_ cells suggests a potential role for this axis in promoting immunological ignorance within HBV-associated HCC TME.

To substantiate these findings, we analyzed a publicly available scRNA-seq dataset (GSE282343) comprising five HBV-HCC and three non-HBV-HCC samples from the NCBI GEO database using our processing pipeline (24). Consistent with our results, the validation cohort demonstrated TIGIT-NECTIN2 interactions involving T_EM_, T_EMRA_, T_PEX_, and T_EX_ CD8⁺ T cells, detected in HBV-HCC samples and, to a lesser extent, in non-HBV-HCC samples. Specifically, TIGIT-NECTIN2 interactions involved T_EM_ and T_EX_ subsets in three of five HBV-HCC cases, T_EMRA_ and T_PEX_ subsets in two cases, and T_EX_ subsets in one of three non-HBV-HCC cases (Fig. 5D). In contrast, TIGIT-NECTIN3 interactions were observed in T_EM_, T_EMRA_, T_PEX_, and T_EX_ CD8⁺ T cells of HBV-HCC cases (Fig. 5E). Collectively, these findings indicate that NECTIN-TIGIT interactions are preferentially engaged in HBV-associated HCC.

### TIGIT-NECTIN3 Axis as a Distinct Immunosuppressive Pathway in HBV-Associated HCC

Building on our cell-cell interaction data, which highlighted TIGIT-NECTIN3 interactions in shaping the immunosuppressive landscape in HBV-associated HCC, we next examined the expression of TIGIT and NECTIN3 in HCC tissues with or without HBV infection. Immunohistochemistry (IHC) staining for NECTIN3 was performed on a cohort of HBV-high HCC samples (n = 22) (HBV viral load >200 IU/ml) and non-B non-C (NBNC) HCC samples (n = 16) (Table S5). Consistent with our scRNA-seq findings, NECTIN3 expression was significantly higher in HBV-infected HCC compared to NBNC HCC (Fig. 6A and 6B). In line with this, our bulk RNA-seq analysis further showed that NECTIN3 expression was positively correlated with HBsAg and HBx transcripts (Fig. 6C and 6D).

**Fig. 6.**
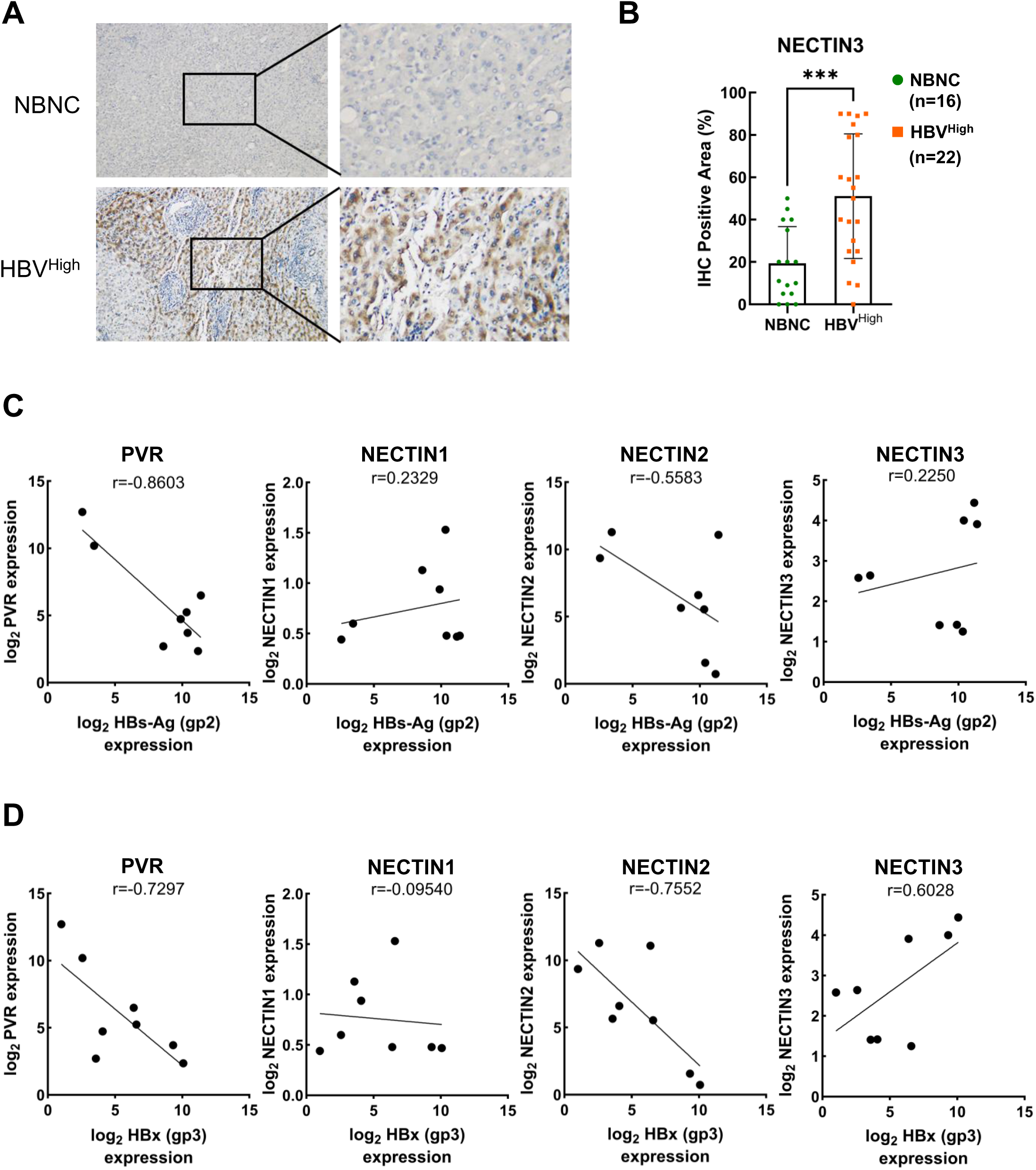
High expression of NECTIN3 in HBV-associated HCC tumor tissues. (A) Representative immunohistochemical (IHC) staining of NECTIN3 in HBV-associated HCC and non-B non-C (NBNC) HCC specimens. NECTIN3-positive staining appears in brown, and nuclei are counterstained with hematoxylin (blue). (B) NECTIN3 expression levels between HBV-associated HCC and NBNC HCC specimens. (C, D) Correlation of HBs-Ag (gp2) (C) and HBx(gp3) (D) expression with PVR, NECTIN1, NECTIN2, and NECTIN3 in HCC tumor tissues.

To assess TIGIT expression within immune subsets, we performed multiplex immunofluorescence staining on the 22 HBV-high and 16 NBNC HCC specimens, evaluating TIGIT levels in CD8⁺ T cells, CD4⁺ T cells, and Treg cells. In line with our single-cell and cell-cell interaction data, higher TIGIT expression levels were observed in both CD8⁺ and Treg cells, the two major T cell populations involved in the NECTIN3-TIGIT interaction, in HBV-high specimens (Fig. 7A). To verify this finding, we performed immune cell deconvolution (35) on bulk RNA-seq data from the TCGA LIHC dataset. Given the high expression of TIGIT in CD8⁺ c4 (Fig. 2E), we examined the proportions of these cell populations in HBV-associated HCC tumors with high and low expression of TIGIT and different NECTIN isoforms. Interestingly, CD8⁺ c4 cells, which are predominantly located at the terminal end of state 3 (Fig. 4F), were significantly more abundant in NECTIN3-high (top 25%, n=95) compared to NECTIN3-low (bottom 25%, n=95) tumors, as well as in TIGIT-high versus TIGIT-low HCCs (Fig. 7B), suggesting the involvement of this specific CD8⁺ T cell subset in the NECTIN3-TIGIT axis. Altogether, our findings reveal a previously unrecognized TIGIT-NECTIN3 axis as a potential HBV-specific immunosuppressive mechanism, highlighting TIGIT-NECTIN3 as a novel mode of crosstalk between T cells and virus-infected HCC cells that may help preserve immune responsiveness and thereby improve sensitivity to ICI therapy.

**Fig. 7.**
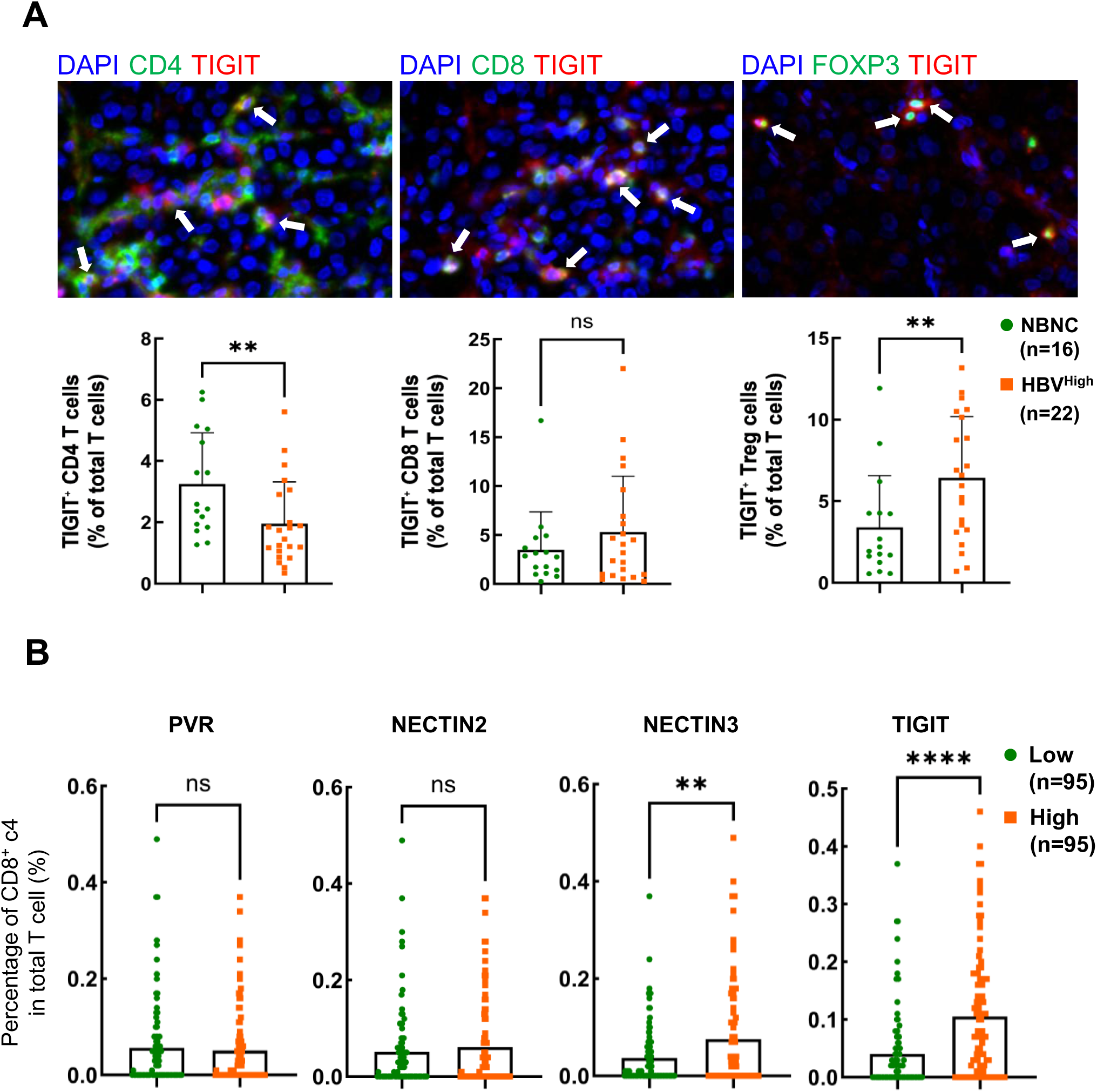
TIGIT–NECTIN expression in CD8⁺ T cells in HBV-associated HCC. (A) Representative multiplex immunofluorescence staining using anti-CD4, anti-CD8, anti-FOXP3, and anti-TIGIT antibodies (upper panel). Quantification of TIGIT-positive cells among CD4⁺ (lower left panel), CD8⁺ (lower middle panel), and FOXP3⁺ (Treg) (lower right panel) T cell populations. (B) Proportion of CD8^+^ c4 T cells in HCC tumors from TCGA LIHC dataset with high (top 25%, n = 95) and low (bottom 25%, n = 95) expression of PVR, NECTIN2, NECTIN3, and TIGIT, as determined by deconvolution analysis of bulk RNA-seq data from TCGA.

## Discussion

In this study, we demonstrate HBV transcripts in scRNA-seq of HCC, enabling direct linkage of viral gene expression and immune modulation at single-cell resolution. We delineate the immunosuppressive landscape, showing that tumor cells engage in distinct TIGIT-NECTIN interactions with CD8⁺ T cells in shaping the immunosuppressive environment in HBV-associated hepatocarcinogenesis. Specifically, TIGIT-NECTIN3 interaction predominates in HCC clusters with high HBV transcripts, whereas TIGIT-NECTIN2 interaction is observed in clusters with high-to-medium HBV transcripts, suggesting a dynamic immunosuppressive mechanism in hepatocarcinogenesis. Furthermore, we identified a novel CD8⁺ T cell population corresponding to the recently proposed ignorant T cells, which may play a critical role in responsiveness to ICIs (16). Importantly, integration of bulk RNA-seq data from tumor and adjacent non-tumor tissues with scRNA-seq-defined CD8⁺ T cell subsets highlighted CD24, STMN1, and EZH2 as potential biomarkers of this ignorant CD8⁺ T cell population in HBV-associated HCC.

Given the pioneering role of virus research in cancer biology and the observed favorable responses to ICIs in virus-associated HCC (14, 15), a deeper understanding of how HBV modulates tumor-immune interactions and shapes immune cells is urgently needed to decipher the mechanisms underlying the differential responses to ICIs among patients. In particular, CD8⁺ T cells, also known as cytotoxic T lymphocytes (CTLs), are the primary immune cells responsible for controlling intracellular viral infections and mediating anti-tumor immune response. However, during chronic infection and tumor progression, CD8⁺ T cells can enter a state of dysfunction known as exhaustion, characterized by sustained expression of inhibitory receptors and diminished effector function (36). Notably, T_PEX_ cells exhibit stem-like and proliferative capacities that enable adaptation to persistent antigen exposure and sustain T cell immunity, underscoring their pivotal role in mediating antitumor responses to immunotherapy.

Importantly, Galluzzi *et al.* recently proposed a ten-step model of disease-targeting T cell immunity (16). In this model, CD8⁺ T cell exhaustion arises at step 8, where persistent antigen exposure combined with immunosuppressive signals in diseased tissue drives the transition from stem-like T_PEX_ cells to T_EX_ cells. At step 9, defective antigen presentation by disease cells leads to immunological ignorance, a state in which CD8⁺ T cells retain a naive-like phenotype (16). While the immune landscape of the TME in HBV-related HCC has been extensively explored, the trajectory of CD8⁺ T cells toward T_PEX_ or immunological ignorance fates remains unknown. In this study, trajectory analysis revealed a T_PEX_-like population emerging after T_EX,_ corresponding to immunologically ignorant CD8⁺ T cells. Unlike T_PEX_ cells, which are defined by PD-1⁺TCF-1⁺TOX⁺ expression (33), CD8⁺ T cells in the immunological ignorance state lack well-established phenotypic markers. By integrating scRNA-seq and bulk RNA-seq data, we identified CD24, STMN1, and EZH2 as potential biomarkers for this immunologically ignorant CD8⁺ T cell population. A major limitation that warrants consideration is the limited scRNA-seq sample size. However, we validated our findings using FFPE staining, bulk RNA-seq, bioinformatic deconvolution of TCGA datasets, and an independent publicly available scRNA-seq dataset (GSE282343) comprising five HBV-HCC and three non-HBV-HCC samples. Identification of the features of ignorant CD8⁺ T cells hold important clinical implications for improving immunotherapeutic strategies.

Addressing the emergence of immunologically ignorant CD8⁺ T cells requires a comprehensive view of the HCC TME, encompassing both tumor and immune compartments. Although single-cell analyses have been widely used to study the TME, most HCC studies have focused on immune subsets, leaving tumor compartments less characterized. Here, we applied CellChat analysis on our scRNA-seq data to construct a detailed interaction landscape between tumor and immune compartments. This approach allowed us to identify key immunological features and their potential interactions with HBV-infected HCC cells, providing critical insight into mechanisms driving the emergence and maintenance of immunologically ignorant CD8⁺ T cells. In line with previous observations that HBV-specific T cell dysfunction does not relate solely to PD-1 expression (37), we found that TIGIT was upregulated and at a higher level in HBV-infected HCC, suggesting that multiple co-inhibitory pathways contribute to T cell dysfunction within the unique hepatic microenvironment shaped by HBV. Notably, we identified TIGIT-NECTIN3 interactions specifically between T_PEX_ CD8⁺ T cells and HCC-c1 and HCC-c2 clusters, which exhibit high viral transcript levels. In contrast, TIGIT-NECTIN2 interactions predominantly occur between T_PEX_ CD8⁺ T cells and HCC-1 and HCC-2 clusters, as well as in HCC-c5 and HCC-6 clusters with intermediate viral transcription, suggesting a shift from TIGIT-NECTIN3 to TIGIT-NECTIN2 engagement during the decline of viral transcription.

In addition to primary poliovirus receptor (PVR), TIGIT also engages NECTIN family proteins, including NECTIN1 (PVRL1), NECTIN2 (PVRL2), NECTIN3 (PVRL3), and NECTIN4 (PVRL4), with varying binding affinities (38, 39), highlighting the added complexity of TIGIT-mediated immune regulation in tumor immunology. Nectins are adhesion molecules that also function as viral receptors (40). Both PVR and NECTIN2 directly bind TIGIT and deliver inhibitory signals to immune cells, although TIGIT binds NECTIN2 with much lower affinity than PVR (41). Beyond direct TIGIT binding, NECTIN1 was shown to stabilize PVR, thereby enhancing the PVR-TIGIT interaction, suppressing cytotoxic T cell responses, and ultimately increasing tumor resistance to anti-PD-1 therapies (42). Similarly, NECTIN4 overexpression was found to increase PVR levels on T cell surfaces (39). In contrast, NECTIN3 binds PVR and promotes its internalization (42). Collectively, these findings suggest that NECTIN3 promotes PVR internalization and potentially shifts TIGIT engagement toward TIGIT-NECTIN2-mediated immune suppression. Intriguingly, our data show that NECTIN3 is upregulated in HBV-enriched HCC cells, potentially shifting TIGIT signaling toward NECTIN2/3 interactions, thereby attenuating TIGIT-mediated inhibition of cytotoxic T cells. Thus, we hypothesize that this could alleviate immune suppression and contribute to the emergence of immunological ignorance in the context of chronic HBV infection.

In conclusion, our findings uncover the elusive immunologically ignorant CD8 T cell in HBV-associated HCC and identify candidate biomarkers indicative of this phenotype. NECTIN3 expression in HBV-related HCC mediates a noncanonical HCC-T cell interaction involving TIGIT-NECTIN signaling, with TIGIT-NECTIN2 interactions predominating as viral transcription declines. This dynamic TIGIT-NECTIN signaling landscape may contribute to the emergence of immunological ignorance of T cells in HBV-related HCC.

## Supplementary information

Supplementary data for this article are available in the supplementary file.

## Acknowledgements

The authors acknowledge the scRNA-seq technical services provided by Kate Hua at the National Genomics Center for Clinical and Biotechnological Applications of the Cancer and Immunology Research Center at National Yang Ming Chiao Tung University, and the National Core Facility for Biopharmaceuticals (NCFB) of the National Science and Technology Council. The authors also acknowledge Chih-Sin Hsu and Kai-Hsiang Hsu for their technical expertise in scRNA-seq data analysis, and Yin-Cheng Chen for bioinformatics figure preparation.

## Author contributions

Pei-Ching Chang, Chun-Ying Wu, and Yan-Hwa Wu Lee served as the conceptual leaders of the study. Zhi-Chen Yan, Yung-Chih Hong, Kai-Ting Yu, Tze-Yun Hu, Chen-Ching Lin, Tai-Ming Ko, and Jinn-Moon Yang developed the methodology. Pei-Ching Chang, Zhi-Chen Yan, Yung-Chih Hong, Kai-Ting Yu, Tze-Yun Hu, Pao-Shu Wu, Muh-Hwa Yang, and Jiunn-Chang Lin conducted the experiments and investigations. Pei-Ching Chang, Muh-Hwa Yang, Chun-Ying Wu, and Yan-Hwa Wu Lee acquired funding for the study. Pei-Ching Chang and Yan-Hwa Wu Lee supervised the project. Pei-Ching Chang, Zhi-Chen Yan, and Yung-Chih Hong wrote the original draft, and Yan-Hwa Wu Lee reviewed and edited the manuscript. All authors provided comments on and approved the final manuscript.

## Data availability

All datasets used and analyzed in this study are provided in the supplementary materials. Raw data are available from the corresponding author upon reasonable request.

## Declaration

### Ethical approval

This study was approved by the Institutional Review Board (IRB) of MacKay Memorial Hospital, Taiwan (Approval No. 13MMHIS269).

