## Supplementary Figures for "TIGIT-NECTIN2/3 signaling preserves ignorant CD8⁺ T cells for favorable immune checkpoint outcomes in HBV-related hepatocellular carcinoma"

### Supplementary Methods

#### Single-cell RNA sequencing (scRNA-seq)

Surgically resected HCC tissues were immediately immersed in MACS Tissue Storage Solution (Miltenyi Biotec) and transported on ice to the laboratory. Single-cell suspensions were prepared using Human Tumor Dissociation Kit (Miltenyi Biotec). Tissue blocks were minced into ~1 mm cubes and transferred to gentleMACS C tubes (Miltenyi Biotec) containing digestion enzymes diluted in 5 mL Opti-MEM. Dissociation was performed at 37°C using the gentleMACS Octo Dissociator (Miltenyi Biotec) with the h_TDK_2 program. The suspension was diluted with 5 mL Opti-MEM and filtered through a 70 μm cell strainer (FALCON). Viable cells were enriched via Ficoll density gradient centrifugation (1,000 × g, 15 min, room temperature), and viability was assessed using the Countess II FL Automated Cell Counter (Invitrogen). Samples with viability >90% were centrifuged (300 × g, 7 min, 4°C) and resuspended in Opti-MEM at a density of 1,000 cells/μL. A total of 8,000 single cells per sample were encapsulated by droplet-based microfluidics technology with 10x Genomics Single-Cell-A Chip (10x Genomics). Libraries were prepared using the Chromium Next GEM Single Cell 3′ Reagent Kits v3.1 (10x Genomics) according to the manufacturer's instructions. Barcoded RNAs were pooled and sequenced on an Illumina NovaSeq 6000 platform (paired-end 150 bp, ~150 million reads per sample) at the National Genomics Center for Clinical and Biotechnological Applications of the Cancer and Immunology Research Center, National Yang Ming Chiao Tung University.

Initial analysis was performed using the Cell Ranger pipeline (10x Genomics, v7.0.0) for demultiplexing, barcode processing, and FASTQ file generation. Raw reads were aligned to the human reference genome (GRCh38) to generate alignment matrix files. Reads passing quality control were processed with CellRanger count to obtain unique molecular identifier (UMI) matrices. Barcodes with low UMI counts, low feature counts, or high mitochondrial gene content were filtered out. Doublets were identified and removed using DoubletFinder [1], and the resulting singlet cell counts are summarized in Table S1. Downstream analysis was performed in Partek Flow (Partek Inc.), including $\log_2$(CPM+1) normalization, noise reduction by removal of genes expressed in fewer than 1% of cells, batch effect correction using the Seurat v3 integration method [2], and dimensionality reduction via principal component analysis (PCA). Using PCA with 11 components and a resolution of 0.5, we identified 13 cell clusters via unsupervised graph-based clustering and visualized the results using t-SNE. Cell types were initially assigned by CIRP, a web-based R/shiny app and R package to annotate cell clusters in single cell RNA sequencing experiments, [3] and further refined based on canonical markers from Sun *et al.* [4]. For subset analyses, T cell clustering was performed using 8 PCA components and a resolution of 0.5, while HCC cell clustering used 12 PCA components with the same resolution. The specific T-cell subsets identify by cell identities were annotated by reference [5, 6] mapping using the ProjecTILs R package [7].

#### Bulk RNA sequencing (RNA-seq)

HCC tissues were homogenized in 1 ml of TRIzol reagent (Invitrogen, 15596-018) by Precellys® Tissue Homogenizing Mixed Beads Kit (Bertin) using four 15-second cycles of homogenization at 6,500 rpm. The homogenates were centrifuged at 12,000 g for 10 min at 4°C. The supernatants were collected, and total RNA was isolated. RNA-seq libraries were prepared by Illumina® Stranded mRNA Prep, ligation kit (Illumina, 20040534) using 4 µg of total RNA, and rRNA was depleted using the QIAseq FastSelect-rRNA HMR kit (Qiagen, 334386) according to the manufacturer's instructions. High-throughput RNA sequencing (2*150bp) was performed by the sequencing core facility of the Cancer and Immunology Research Center at National Yang Ming Chiao Tung University using the NovaSeq X plus (Illumina). The raw reads were aligned to the human genome GRCh38/Hg38 using CLC genomics Workbench v21.0.5 (Qiagen). Transcript levels were expressed as reads per kilobase of transcript per million mapped reads (RPKM) with mRNA information obtained from RefSeq using PartekFlow (Partek Inc., USA). Differential mRNA expression was analyzed using RPKM values and expressed as fold change.

#### Multiplex immunofluorescence staining

CD3 (Abcam), CD4 (Invitrogen), CD8 (Invitrogen), FOXP3 (Abcam), and TIGIT (Abcam) antibodies were subsequently applied, followed by Opal Polymer HRP Ms+Rb incubation and Opal fluorophore conjugated tyramide signal amplification. The slides were autoclave heat-treated after each signal amplification step to remove bound antibodies. Nuclei were stained with DAPI after antigens were labeled. The slides were then scanned to acquire multispectral images using a Vectra® Polaris™ Automated Quantitative Pathology Imaging system (PerkinElmer). For each slide, 25 fields were selected by a Phenochart slide viewer (Akoya Biosciences) and quantified using InForm image analysis software (Akoya Biosciences).

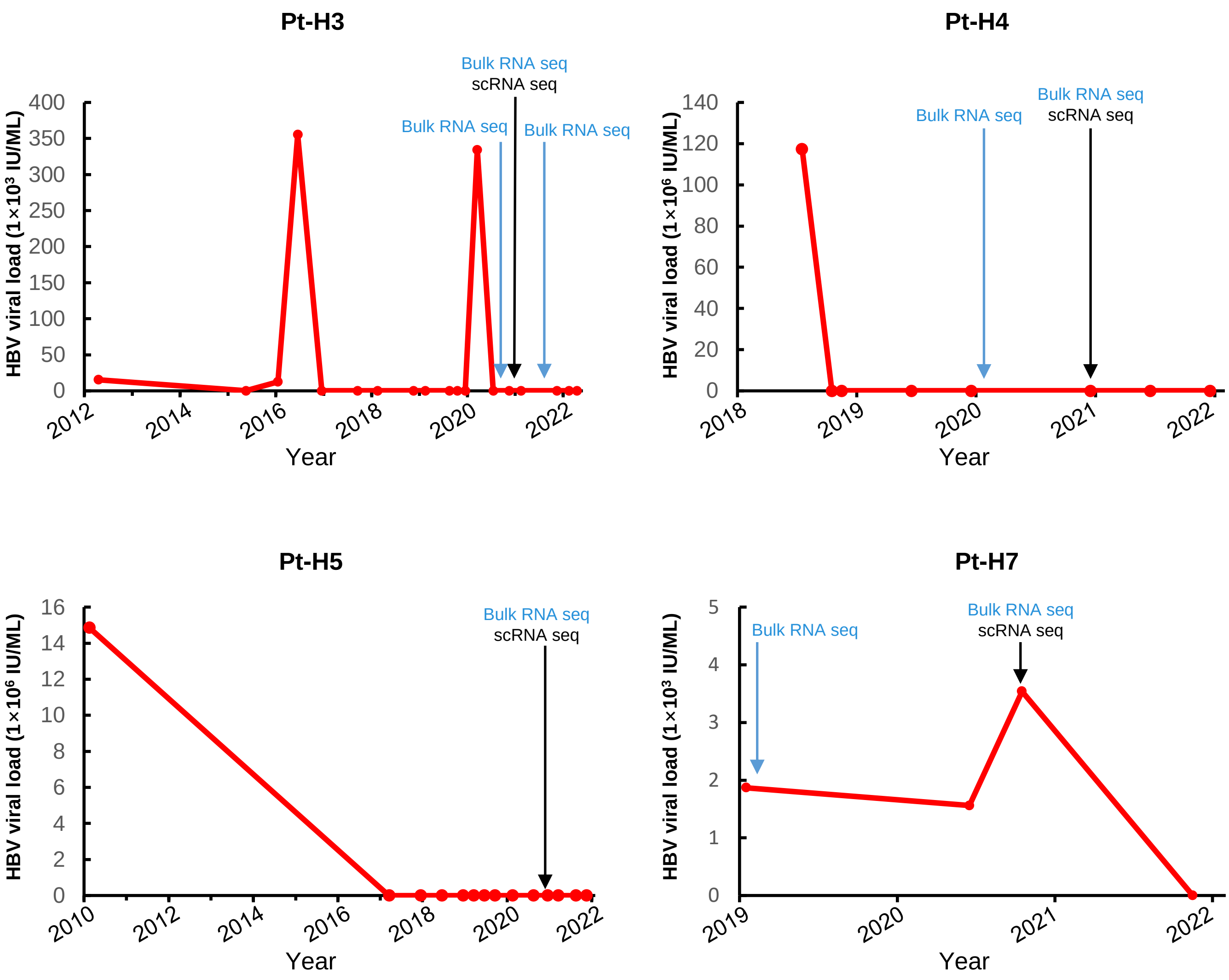


**Figure S1.** Serum HBV DNA levels in patients with HBV-associated hepatocellular carcinoma (HCC).

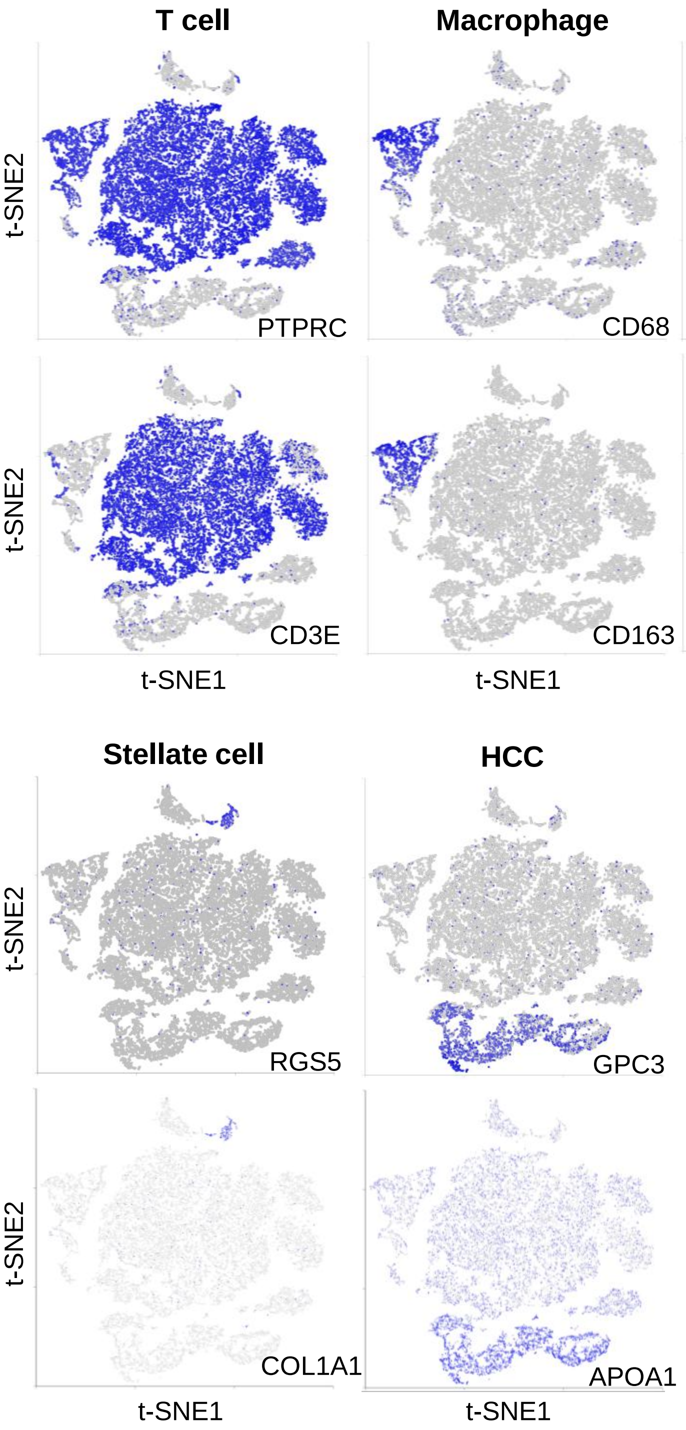


| Cell type | Marker |
|---|---|
| Pan-leukocyte | PTPRC |
| Pan-T cell | CD3E |
| Pan-macrophage | CD68 & CD163 |
| B cell | CD19 & CD79A |
| Endothelial cell | CDH5 & FLT1 |
| Stellate cell | RGS5 & COL1A1 |
| HCC cell | GPC3 & APOA1 |

**Figure S2.** Marker genes used for the identification of major cell types.

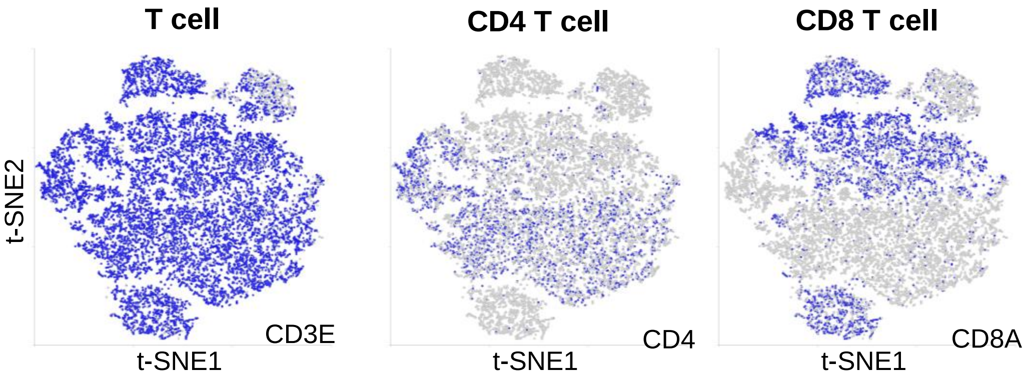


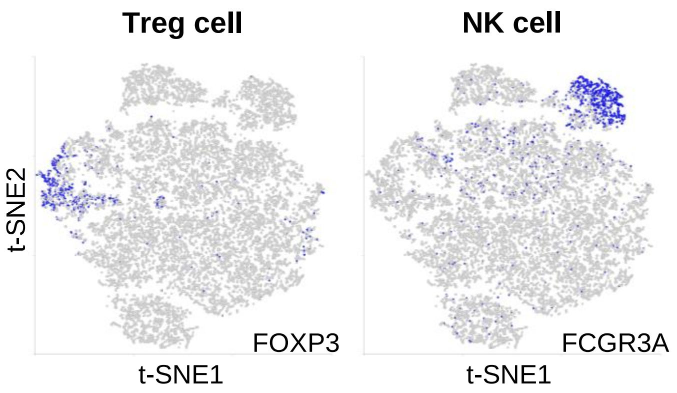


| Cell type | Marker |
|---|---|
| Pan-T cell | CD3E |
| CD4 T cell | CD4 |
| CD8 T cell | CD8A |
| Treg cell | FOXP3 |
| NK cell | FCGR3A |

**Figure S3.** Marker genes used for the identification of immune cell subsets.

Table S1. Pathological data of the HBV-associated HCC patients

| **Case number** | **Pt-H3** | **Pt-H4** | **Pt-H5** | **Pt-H7** |
|---|---|---|---|---|
| Gender | Male | Male | Male | Female |
| Age | 71 | 44 | 68 | 41 |
| No. of tumor nodules | 1 | 1 | 1 | 1 |
| Tumor size (cm) | 3.2 | 3.5 | 3.5 | 1 |
| Tumor encapsulation | complete | none | partial | complete |
| Venous invasion | - | - | - | - |
| Tumor grade | Ⅱ | Ⅱ | Ⅱ | Ⅱ |
| Background liver | Chronic hepatitis | Liver cirrhosis | Fatty liver | Chronic hepatitis |
| BCLC | A | A | A | 0 |
| HBV status | + | + | + | + |
| HCV status | - | - | - | - |
| HDV status | - | - | - | - |
| HBeAg | NA | NA | NA | NA |
| Anti-HBc | - | - | - | - |
| HBV DNA (IU/ml) | <20 | 17.2 | <20 | 3540 |
| Initial Diagnose<br>Recurrent | 1st (20200820)<br>**2nd (20201207)**<br>3rd (20211007) | 1st (20180605)<br>2nd (20181001)<br>3rd (20200109)<br>**4th (20201217)** | **1st (20201221)** | 1st (20190221)<br>**2nd (20201231)** |

BCLC: Barcelona Clinic Liver Cancer Classification; +: positive; -: negative; NA: none available
Bold: scRNA-seq

**Table S1**

Table S2. Summary of the scRNA-seq datasets

| | Pt-H3 | Pt-H4 | Pt-H5 | Pt-H7 |
|---|---|---|---|---|
| Raw cell counts | 7,190 | 6,081 | 5,526 | 9,291 |
| UMI counts | >1,800 | >2,000 | >1,800 | >2,000 |
| Cell counts after UMI | 6,514 | 5,425 | 4,835 | 8,478 |
| Feature counts | <1,000 | <800 | <1,000 | <1,000 |
| Cell counts after Feature | 6,336 | 5,334 | 4,700 | 8,007 |
| Mitochondrial gene | <20% | <20% | <20% | <20% |
| Cell counts after Mitochondria gene | 6,197 | 4,781 | 4,669 | 7,997 |
| Doublet | 307 | 183 | 174 | 512 |
| Singlet cell counts | 5,890 | 4,598 | 4,495 | 7,485 |



**Table S2**

Table S3. Summary of ProjecTILs-Defined $CD8^+$ T-Cell Subtypes Across Pseudotime Trajectory States.

| | State (%) | | | | |
|---|---|---|---|---|---|
| CD8 subtypes | 1 | 2 | 3 | 4 | 5 |
| CD8 Naïve-like | 68.7 | 1.4 | 28.5 | 0.1 | 1.3 |
| CD8 $T_{CM}$ | 25.4 | 7.0 | 29.1 | 11.8 | 26.7 |
| CD8 $T_{EM}$ | 0.3 | 2.2 | 40.1 | 19.6 | 37.8 |
| CD8 $T_{EMRA}$ | 6.5 | 7.6 | 59.8 | 12.0 | 14.1 |
| CD8 $T_{PEX}$ | 2.3 | 0.0 | 97.7 | 0.0 | 0.0 |
| CD8 $T_{EX}$ | 0.4 | 1.2 | 86.1 | 7.0 | 5.4 |
| CD8 $T_{MAIT}$ | 0.2 | 0.8 | 27.8 | 20.6 | 50.5 |

Table S4. CD8$^+$ T-cell c4 cluster-specific biomarkers

| | Non-Tumor | | | | | | | | **Tumor** | | | | | | | | | |
|---|---|---|---|---|---|---|---|---|---|---|---|---|---|---|---|---|---|---|
| CD8$^+$ C4 Biomarker | Pt-H3-1 | Pt-H3 | Pt-H3-3 | Pt-H4-3 | Pt-H4 | Pt-H5 | Pt-H7-1 | Pt-H7 | **Pt-H3-1** | **Pt-H3** | **Pt-H3-3** | **Pt-H4-3** | **Pt-H4** | **Pt-H5** | **Pt-H7-1** | **Pt-H7** | **Non-Tumor Average** | **Tumor Average** |
| CD24 | 0.92 | 2.00 | 0.81 | 2.46 | 1.86 | 0.34 | 4.07 | 0.56 | **52.70** | **33.85** | **35.16** | **0.06** | **0.73** | **5.68** | **41.47** | **24.18** | 1.63 | **24.23** |
| HIST1H3B | 0.16 | 0.21 | 0.17 | 0.12 | 1.10 | 0.05 | 1.11 | 0.51 | **9.95** | **6.35** | **3.37** | **25.64** | **26.60** | **11.17** | **8.39** | **5.96** | 0.43 | **12.18** |
| MKI67 | 0.10 | 0.12 | 0.09 | 0.18 | 0.99 | 0.15 | 0.49 | 0.15 | **6.23** | **4.54** | **1.54** | **17.66** | **13.26** | **7.32** | **2.77** | **2.00** | 0.28 | **6.92** |
| ASPM | 0.15 | 0.18 | 0.08 | 0.33 | 0.84 | 0.20 | 0.54 | 0.10 | **9.35** | **5.87** | **2.10** | **13.16** | **6.66** | **6.62** | **4.06** | **2.41** | 0.30 | **6.28** |
| HIST1H1B | 0.16 | 0.06 | 0.10 | 0.14 | 1.20 | 0.21 | 0.99 | 0.33 | **3.35** | **1.98** | **1.45** | **17.13** | **13.62** | **7.70** | **1.97** | **1.36** | 0.40 | **6.07** |
| NUSAP1 | 0.32 | 0.55 | 0.23 | 0.49 | 1.09 | 0.39 | 1.18 | 0.53 | **4.53** | **3.35** | **1.76** | **13.20** | **12.03** | **5.56** | **3.29** | **2.66** | 0.60 | **5.80** |
| TOP2A | 0.03 | 0.06 | 0.04 | 0.07 | 0.54 | 0.04 | 0.24 | 0.05 | **4.15** | **2.61** | **2.81** | **9.91** | **7.74** | **4.12** | **2.30** | **1.79** | 0.13 | **4.43** |
| TPX2 | 0.16 | 0.14 | 0.10 | 0.16 | 0.53 | 0.14 | 0.45 | 0.17 | **3.58** | **1.80** | **1.87** | **9.67** | **8.17** | **3.83** | **1.75** | **1.23** | 0.23 | **3.99** |
| STMN1 | 0.32 | 0.36 | 0.32 | 0.28 | 0.69 | 0.28 | 0.88 | 0.22 | **1.76** | **1.08** | **1.87** | **10.14** | **9.73** | **2.87** | **0.97** | **1.01** | 0.42 | **3.68** |
| EZH2 | 0.31 | 0.31 | 0.20 | 0.29 | 0.67 | 0.28 | 0.55 | 0.21 | **2.71** | **2.00** | **1.38** | **7.14** | **4.90** | **4.08** | **1.27** | **1.22** | 0.35 | **3.09** |
| MCM4 | 0.30 | 0.36 | 0.36 | 0.28 | 0.56 | 0.37 | 0.50 | 0.33 | **3.14** | **2.07** | **1.02** | **3.60** | **4.16** | **5.58** | **2.11** | **1.56** | 0.38 | **2.90** |
| CXCL13 | 0.16 | 0.00 | 0.09 | 0.00 | 0.00 | 0.00 | 0.10 | 0.12 | **0.42** | **0.90** | **0.40** | **0.03** | **0.52** | **0.28** | **8.32** | **11.64** | 0.06 | **2.81** |
| TYMS | 0.10 | 0.10 | 0.15 | 0.17 | 0.26 | 0.19 | 0.19 | 0.14 | **1.72** | **0.96** | **0.55** | **4.82** | **5.87** | **1.95** | **1.65** | **1.39** | 0.16 | **2.37** |
| ZWINT | 0.03 | 0.09 | 0.01 | 0.04 | 0.43 | 0.11 | 0.29 | 0.06 | **1.62** | **0.89** | **0.72** | **6.70** | **5.72** | **1.68** | **0.76** | **0.56** | 0.13 | **2.33** |
| ECT2 | 0.11 | 0.12 | 0.09 | 0.13 | 0.34 | 0.10 | 0.40 | 0.11 | **2.19** | **1.60** | **1.51** | **4.08** | **3.82** | **2.19** | **1.52** | **1.19** | 0.18 | **2.26** |
| FAM111B | 0.02 | 0.01 | 0.02 | 0.11 | 0.31 | 0.02 | 0.16 | 0.03 | **1.24** | **1.09** | **1.00** | **6.48** | **3.55** | **1.54** | **1.34** | **1.24** | 0.09 | **2.19** |
| UBE2C | 0.03 | 0.03 | 0.08 | 0.08 | 0.27 | 0.07 | 0.15 | 0.04 | **2.71** | **1.64** | **1.02** | **3.90** | **4.22** | **2.19** | **0.95** | **0.65** | 0.09 | **2.16** |
| CDKN3 | 0.01 | 0.02 | 0.00 | 0.00 | 0.20 | 0.06 | 0.16 | 0.06 | **1.23** | **0.92** | **1.38** | **3.71** | **5.47** | **1.50** | **1.68** | **1.10** | 0.06 | **2.12** |
| CD109 | 0.13 | 0.18 | 0.10 | 0.13 | 0.41 | 0.10 | 0.40 | 0.09 | **0.19** | **0.16** | **1.27** | **5.96** | **5.29** | **3.54** | **0.30** | **0.25** | 0.19 | **2.12** |
| CENPF | 0.03 | 0.04 | 0.01 | 0.04 | 0.13 | 0.05 | 0.14 | 0.04 | **3.36** | **1.77** | **1.16** | **2.81** | **1.71** | **4.15** | **1.05** | **0.84** | 0.06 | **2.11** |
| DTL | 0.03 | 0.05 | 0.05 | 0.06 | 0.21 | 0.06 | 0.14 | 0.06 | **2.14** | **1.23** | **0.85** | **4.00** | **3.13** | **2.62** | **1.35** | **0.97** | 0.08 | **2.03** |
| CCNB1 | 0.08 | 0.08 | 0.07 | 0.10 | 0.13 | 0.11 | 0.18 | 0.11 | **2.74** | **1.70** | **0.84** | **2.93** | **3.38** | **2.43** | **1.02** | **0.84** | 0.11 | **1.99** |
| KNL1 | 0.15 | 0.23 | 0.20 | 0.27 | 0.37 | 0.22 | 0.37 | 0.13 | **1.98** | **1.46** | **0.56** | **4.13** | **3.25** | **1.56** | **1.32** | **1.19** | 0.24 | **1.93** |
| FOXM1 | 0.04 | 0.03 | 0.01 | 0.05 | 0.19 | 0.04 | 0.15 | 0.04 | **2.59** | **1.27** | **1.23** | **2.52** | **1.98** | **3.38** | **1.50** | **0.96** | 0.07 | **1.93** |
| NUF2 | 0.01 | 0.01 | 0.02 | 0.01 | 0.43 | 0.01 | 0.10 | 0.02 | **0.83** | **0.52** | **0.51** | **6.31** | **4.91** | **1.19** | **0.49** | **0.37** | 0.08 | **1.89** |
| HELLS | 0.09 | 0.12 | 0.07 | 0.16 | 0.27 | 0.12 | 0.25 | 0.10 | **2.72** | **1.54** | **0.40** | **3.40** | **2.42** | **1.83** | **1.04** | **0.76** | 0.15 | **1.76** |
| E2F1 | 0.03 | 0.09 | 0.02 | 0.07 | 0.20 | 0.09 | 0.17 | 0.04 | **1.60** | **1.17** | **0.47** | **3.25** | **1.96** | **2.46** | **1.52** | **1.66** | 0.09 | **1.76** |
| MYBL2 | 0.02 | 0.02 | 0.01 | 0.03 | 0.16 | 0.02 | 0.04 | 0.02 | **3.22** | **1.40** | **0.50** | **2.85** | **2.00** | **2.41** | **0.50** | **0.46** | 0.04 | **1.67** |
| ANLN | 0.02 | 0.03 | 0.01 | 0.03 | 0.19 | 0.03 | 0.13 | 0.03 | **1.66** | **0.96** | **0.73** | **3.69** | **2.73** | **2.13** | **0.74** | **0.65** | 0.06 | **1.66** |
| NCAPG2 | 0.24 | 0.28 | 0.29 | 0.30 | 0.48 | 0.37 | 0.34 | 0.24 | **1.18** | **0.79** | **1.11** | **3.76** | **2.55** | **2.01** | **0.74** | **0.76** | 0.32 | **1.61** |
| AURKA | 0.20 | 0.16 | 0.16 | 0.13 | 0.30 | 0.18 | 0.21 | 0.09 | **1.06** | **0.63** | **2.25** | **1.79** | **3.86** | **1.00** | **1.48** | **0.68** | 0.18 | **1.59** |
| TK1 | 0.04 | 0.07 | 0.07 | 0.08 | 0.16 | 0.08 | 0.19 | 0.14 | **0.92** | **0.56** | **0.35** | **3.42** | **3.02** | **0.60** | **1.09** | **0.74** | 0.10 | **1.34** |
| FANCI | 0.14 | 0.17 | 0.09 | 0.14 | 0.33 | 0.15 | 0.24 | 0.14 | **1.27** | **0.71** | **0.52** | **2.99** | **2.60** | **1.00** | **0.64** | **0.51** | 0.17 | **1.28** |

Table S4. CD8⁺ T-cell c4 cluster-specific biomarkers (continue)

| | Non-Tumor | | | | | | | | **Tumor** | | | | | | | | | |
|---|---|---|---|---|---|---|---|---|---|---|---|---|---|---|---|---|---|---|
| $CD8^{+}$ C4 Biomarker | Pt-H3-1 | Pt-H3 | Pt-H3-3 | Pt-H4-3 | Pt-H4 | Pt-H5 | Pt-H7-1 | Pt-H7 | **Pt-H3-1** | **Pt-H3** | **Pt-H3-3** | **Pt-H4-3** | **Pt-H4** | **Pt-H5** | **Pt-H7-1** | **Pt-H7** | **Non-Tumor Average** | **Tumor Average** |
| CDK1 | 0.01 | 0.02 | 0.01 | 0.06 | 0.28 | 0.01 | 0.11 | 0.02 | **0.91** | **0.47** | **0.43** | **3.13** | **3.77** | **0.71** | **0.48** | **0.31** | 0.07 | **1.28** |
| MELK | 0.01 | 0.02 | 0.01 | 0.02 | 0.11 | 0.03 | 0.08 | 0.01 | **0.89** | **0.72** | **0.40** | **2.41** | **2.55** | **1.33** | **0.61** | **0.57** | 0.03 | **1.18** |
| SPC24 | 0.02 | 0.01 | 0.01 | 0.03 | 0.22 | 0.04 | 0.08 | 0.05 | **1.34** | **1.01** | **0.28** | **2.64** | **1.67** | **0.81** | **0.60** | **0.47** | 0.06 | **1.10** |
| NCAPG | 0.02 | 0.00 | 0.01 | 0.02 | 0.13 | 0.03 | 0.08 | 0.03 | **1.16** | **0.67** | **0.72** | **2.26** | **1.88** | **1.13** | **0.49** | **0.39** | 0.04 | **1.09** |
| MCM2 | 0.06 | 0.06 | 0.05 | 0.08 | 0.11 | 0.06 | 0.15 | 0.06 | **1.15** | **0.64** | **0.57** | **1.96** | **1.82** | **1.41** | **0.66** | **0.44** | 0.08 | **1.08** |
| AURKB | 0.07 | 0.00 | 0.05 | 0.08 | 0.12 | 0.08 | 0.14 | 0.05 | **1.19** | **0.59** | **0.28** | **2.25** | **1.66** | **1.44** | **0.69** | **0.53** | 0.07 | **1.08** |
| BUB1B | 0.03 | 0.02 | 0.01 | 0.05 | 0.15 | 0.01 | 0.09 | 0.01 | **1.76** | **0.98** | **0.29** | **1.99** | **1.50** | **1.16** | **0.55** | **0.34** | 0.05 | **1.07** |
| TROAP | 0.02 | 0.01 | 0.03 | 0.01 | 0.15 | 0.03 | 0.08 | 0.04 | **1.03** | **0.79** | **0.52** | **2.36** | **1.62** | **1.30** | **0.48** | **0.45** | 0.04 | **1.07** |
| HMMR | 0.05 | 0.13 | 0.08 | 0.11 | 0.18 | 0.09 | 0.10 | 0.08 | **0.54** | **0.39** | **0.41** | **2.96** | **2.72** | **1.04** | **0.25** | **0.13** | 0.10 | **1.06** |
| HJURP | 0.01 | 0.03 | 0.01 | 0.00 | 0.06 | 0.01 | 0.10 | 0.04 | **1.14** | **0.87** | **0.43** | **1.43** | **1.09** | **1.65** | **0.78** | **0.86** | 0.03 | **1.03** |
| PYCR1 | 0.08 | 0.03 | 0.08 | 0.06 | 0.10 | 0.14 | 0.12 | 0.08 | **3.01** | **1.54** | **0.06** | **1.64** | **1.42** | **0.04** | **0.10** | **0.07** | 0.09 | **0.98** |
| DEPDC1 | 0.01 | 0.00 | 0.01 | 0.02 | 0.11 | 0.00 | 0.07 | 0.02 | **0.53** | **0.21** | **0.41** | **3.15** | **2.03** | **1.09** | **0.25** | **0.21** | 0.03 | **0.98** |
| KIF14 | 0.04 | 0.07 | 0.03 | 0.04 | 0.13 | 0.07 | 0.08 | 0.06 | **0.97** | **0.49** | **0.63** | **1.76** | **1.24** | **1.74** | **0.50** | **0.45** | 0.06 | **0.97** |
| SGO1 | 0.05 | 0.07 | 0.06 | 0.08 | 0.12 | 0.03 | 0.13 | 0.05 | **1.04** | **0.53** | **0.35** | **2.38** | **1.82** | **0.80** | **0.51** | **0.27** | 0.08 | **0.96** |
| CDCA8 | 0.05 | 0.04 | 0.04 | 0.06 | 0.13 | 0.07 | 0.08 | 0.02 | **0.84** | **0.41** | **0.40** | **2.32** | **1.72** | **1.46** | **0.22** | **0.32** | 0.06 | **0.96** |
| TTK | 0.02 | 0.00 | 0.01 | 0.04 | 0.14 | 0.02 | 0.06 | 0.02 | **1.07** | **0.61** | **0.38** | **2.09** | **1.79** | **0.86** | **0.47** | **0.32** | 0.04 | **0.95** |
| KIF4A | 0.02 | 0.02 | 0.01 | 0.02 | 0.13 | 0.02 | 0.13 | 0.03 | **0.84** | **0.51** | **0.47** | **1.93** | **1.67** | **1.44** | **0.36** | **0.23** | 0.05 | **0.93** |
| ARHGAP11A | 0.03 | 0.04 | 0.05 | 0.05 | 0.15 | 0.06 | 0.13 | 0.04 | **1.20** | **0.65** | **0.38** | **1.90** | **1.43** | **0.83** | **0.52** | **0.38** | 0.07 | **0.91** |
| CDCA7 | 0.03 | 0.02 | 0.00 | 0.02 | 0.00 | 0.03 | 0.10 | 0.02 | **1.69** | **1.05** | **0.06** | **0.02** | **0.08** | **0.14** | **3.10** | **1.11** | 0.03 | **0.91** |
| PBK | 0.01 | 0.00 | 0.00 | 0.02 | 0.06 | 0.00 | 0.00 | 0.00 | **0.54** | **0.22** | **0.61** | **1.86** | **2.44** | **0.60** | **0.37** | **0.40** | 0.01 | **0.88** |
| SKA1 | 0.00 | 0.00 | 0.01 | 0.01 | 0.10 | 0.00 | 0.06 | 0.01 | **0.56** | **0.40** | **0.26** | **2.14** | **2.45** | **0.81** | **0.25** | **0.17** | 0.02 | **0.88** |
| TESC | 0.09 | 0.20 | 0.07 | 0.18 | 0.22 | 0.13 | 0.28 | 0.05 | **0.90** | **0.95** | **0.13** | **0.08** | **0.07** | **0.46** | **3.76** | **0.66** | 0.15 | **0.88** |
| CENPW | 0.00 | 0.11 | 0.09 | 0.12 | 0.16 | 0.05 | 0.02 | 0.10 | **0.88** | **0.74** | **0.55** | **1.22** | **0.97** | **1.19** | **0.70** | **0.75** | 0.08 | **0.88** |
| UCK2 | 0.13 | 0.14 | 0.20 | 0.13 | 0.17 | 0.17 | 0.24 | 0.10 | **2.06** | **1.26** | **0.38** | **0.46** | **0.45** | **1.36** | **0.73** | **0.28** | 0.16 | **0.87** |
| BUB1 | 0.02 | 0.01 | 0.01 | 0.05 | 0.10 | 0.02 | 0.06 | 0.02 | **1.27** | **0.88** | **0.51** | **1.22** | **1.12** | **1.17** | **0.41** | **0.34** | 0.04 | **0.86** |
| CDC20 | 0.00 | 0.02 | 0.01 | 0.00 | 0.07 | 0.01 | 0.03 | 0.01 | **0.72** | **0.43** | **0.40** | **2.18** | **1.60** | **1.12** | **0.25** | **0.20** | 0.02 | **0.86** |
| UHRF1 | 0.02 | 0.02 | 0.00 | 0.07 | 0.11 | 0.02 | 0.09 | 0.02 | **1.41** | **0.90** | **0.17** | **1.24** | **1.00** | **0.81** | **0.51** | **0.40** | 0.04 | **0.80** |
| KIF15 | 0.00 | 0.04 | 0.02 | 0.01 | 0.12 | 0.03 | 0.09 | 0.01 | **0.78** | **0.43** | **0.27** | **2.13** | **1.75** | **0.55** | **0.27** | **0.20** | 0.04 | **0.80** |
| WDHD1 | 0.08 | 0.08 | 0.05 | 0.08 | 0.12 | 0.05 | 0.10 | 0.05 | **1.16** | **0.78** | **0.42** | **1.22** | **0.97** | **1.02** | **0.43** | **0.33** | 0.08 | **0.79** |
| RAD51AP1 | 0.01 | 0.10 | 0.04 | 0.04 | 0.06 | 0.03 | 0.09 | 0.04 | **1.25** | **0.62** | **0.30** | **1.19** | **0.98** | **1.05** | **0.36** | **0.33** | 0.05 | **0.76** |
| CKAP2L | 0.01 | 0.02 | 0.03 | 0.04 | 0.08 | 0.01 | 0.06 | 0.02 | **0.69** | **0.47** | **0.57** | **1.43** | **1.23** | **0.89** | **0.43** | **0.35** | 0.03 | **0.76** |
| CCNB2 | 0.02 | 0.04 | 0.01 | 0.05 | 0.08 | 0.01 | 0.13 | 0.01 | **1.19** | **0.61** | **0.50** | **1.24** | **1.30** | **0.56** | **0.33** | **0.29** | 0.04 | **0.75** |

Table S4. CD8$^+$ T-cell c4 cluster-specific biomarkers (continue)

| | Non-Tumor | | | | | | | | **Tumor** | | | | | | | | | |
|---|---|---|---|---|---|---|---|---|---|---|---|---|---|---|---|---|---|---|
| CD8$^+$ C4 Biomarker | Pt-H3-1 | Pt-H3 | Pt-H3-3 | Pt-H4-3 | Pt-H4 | Pt-H5 | Pt-H7-1 | Pt-H7 | **Pt-H3-1** | **Pt-H3** | **Pt-H3-3** | **Pt-H4-3** | **Pt-H4** | **Pt-H5** | **Pt-H7-1** | **Pt-H7** | **Non-Tumor Average** | **Tumor Average** |
| KIF23 | 0.01 | 0.00 | 0.01 | 0.04 | 0.08 | 0.02 | 0.07 | 0.02 | **0.82** | **0.55** | **0.31** | **1.50** | **1.48** | **0.63** | **0.38** | **0.24** | 0.03 | **0.74** |
| CDT1 | 0.00 | 0.01 | 0.01 | 0.01 | 0.11 | 0.02 | 0.05 | 0.01 | **0.37** | **0.21** | **0.14** | **2.26** | **1.69** | **0.58** | **0.24** | **0.11** | 0.03 | **0.70** |
| DLGAP5 | 0.00 | 0.00 | 0.00 | 0.05 | 0.08 | 0.01 | 0.03 | 0.04 | **0.88** | **0.49** | **0.28** | **1.30** | **1.19** | **0.79** | **0.34** | **0.25** | 0.03 | **0.69** |
| PAQR4 | 0.04 | 0.07 | 0.04 | 0.04 | 0.05 | 0.04 | 0.07 | 0.02 | **0.24** | **0.25** | **0.29** | **1.30** | **1.22** | **1.59** | **0.23** | **0.22** | 0.05 | **0.67** |
| GTSE1 | 0.01 | 0.03 | 0.01 | 0.01 | 0.05 | 0.01 | 0.06 | 0.02 | **1.01** | **0.53** | **0.17** | **1.42** | **0.78** | **0.74** | **0.34** | **0.26** | 0.02 | **0.66** |
| PLK1 | 0.03 | 0.08 | 0.09 | 0.07 | 0.10 | 0.10 | 0.17 | 0.03 | **1.63** | **0.56** | **0.21** | **0.89** | **0.78** | **0.86** | **0.21** | **0.08** | 0.08 | **0.65** |
| BIRC5 | 0.02 | 0.01 | 0.05 | 0.02 | 0.16 | 0.03 | 0.04 | 0.03 | **0.75** | **0.31** | **0.18** | **1.73** | **1.23** | **0.45** | **0.26** | **0.16** | 0.05 | **0.63** |
| CENPE | 0.03 | 0.06 | 0.02 | 0.04 | 0.07 | 0.05 | 0.09 | 0.05 | **1.07** | **0.54** | **0.20** | **1.17** | **0.78** | **0.78** | **0.26** | **0.23** | 0.05 | **0.63** |
| MCM10 | 0.02 | 0.00 | 0.03 | 0.02 | 0.13 | 0.01 | 0.02 | 0.01 | **0.55** | **0.24** | **0.12** | **1.65** | **1.45** | **0.41** | **0.17** | **0.19** | 0.03 | **0.60** |
| NEIL3 | 0.02 | 0.02 | 0.03 | 0.03 | 0.09 | 0.01 | 0.18 | 0.03 | **0.52** | **0.28** | **0.27** | **1.29** | **0.99** | **0.87** | **0.19** | **0.14** | 0.05 | **0.57** |
| CDCA2 | 0.03 | 0.01 | 0.01 | 0.03 | 0.07 | 0.01 | 0.02 | 0.01 | **0.46** | **0.29** | **0.23** | **1.46** | **1.25** | **0.47** | **0.17** | **0.10** | 0.02 | **0.55** |
| CEP55 | 0.01 | 0.02 | 0.01 | 0.03 | 0.04 | 0.00 | 0.04 | 0.03 | **0.36** | **0.32** | **0.22** | **0.95** | **0.80** | **1.03** | **0.21** | **0.22** | 0.02 | **0.51** |
| PCLAF | 0.08 | 0.09 | 0.08 | 0.04 | 0.13 | 0.11 | 0.07 | 0.04 | **0.51** | **0.54** | **0.22** | **0.93** | **0.79** | **0.40** | **0.22** | **0.19** | 0.08 | **0.47** |
| KIF2C | 0.01 | 0.02 | 0.00 | 0.02 | 0.12 | 0.01 | 0.03 | 0.00 | **0.42** | **0.22** | **0.16** | **1.32** | **0.87** | **0.59** | **0.11** | **0.09** | 0.03 | **0.47** |
| CLSPN | 0.00 | 0.01 | 0.02 | 0.01 | 0.06 | 0.04 | 0.07 | 0.02 | **0.33** | **0.18** | **0.16** | **0.92** | **0.76** | **1.01** | **0.18** | **0.11** | 0.03 | **0.46** |
| CHEK1 | 0.05 | 0.07 | 0.02 | 0.03 | 0.06 | 0.04 | 0.11 | 0.03 | **0.72** | **0.38** | **0.09** | **0.74** | **0.72** | **0.52** | **0.30** | **0.16** | 0.05 | **0.45** |
| CDCA5 | 0.01 | 0.02 | 0.00 | 0.03 | 0.07 | 0.01 | 0.05 | 0.01 | **0.67** | **0.31** | **0.27** | **0.91** | **0.51** | **0.46** | **0.32** | **0.19** | 0.03 | **0.45** |
| GINS2 | 0.01 | 0.06 | 0.03 | 0.04 | 0.08 | 0.07 | 0.04 | 0.04 | **0.16** | **0.12** | **0.09** | **1.14** | **1.16** | **0.42** | **0.21** | **0.21** | 0.05 | **0.44** |
| ORC6 | 0.03 | 0.01 | 0.02 | 0.03 | 0.08 | 0.05 | 0.07 | 0.03 | **0.39** | **0.24** | **0.11** | **1.12** | **0.93** | **0.43** | **0.15** | **0.12** | 0.04 | **0.44** |
| CTHRC1 | 0.00 | 0.01 | 0.00 | 0.00 | 0.00 | 0.00 | 0.03 | 0.00 | **1.05** | **0.86** | **0.57** | **0.03** | **0.28** | **0.32** | **0.10** | **0.01** | 0.01 | **0.40** |
| CDC45 | 0.00 | 0.00 | 0.01 | 0.00 | 0.05 | 0.01 | 0.02 | 0.03 | **0.40** | **0.29** | **0.11** | **0.53** | **0.43** | **0.63** | **0.24** | **0.19** | 0.01 | **0.35** |
| CENPA | 0.01 | 0.00 | 0.00 | 0.05 | 0.04 | 0.03 | 0.03 | 0.01 | **0.37** | **0.26** | **0.03** | **0.72** | **0.73** | **0.44** | **0.14** | **0.06** | 0.02 | **0.34** |
| MND1 | 0.03 | 0.00 | 0.01 | 0.00 | 0.04 | 0.00 | 0.01 | 0.00 | **0.31** | **0.13** | **0.30** | **0.41** | **0.37** | **0.30** | **0.22** | **0.24** | 0.01 | **0.28** |
| CDCA3 | 0.02 | 0.03 | 0.02 | 0.01 | 0.05 | 0.03 | 0.05 | 0.00 | **0.30** | **0.18** | **0.11** | **0.58** | **0.39** | **0.35** | **0.12** | **0.12** | 0.03 | **0.27** |
| PKMYT1 | 0.01 | 0.02 | 0.01 | 0.01 | 0.05 | 0.01 | 0.04 | 0.01 | **0.11** | **0.09** | **0.11** | **0.82** | **0.56** | **0.26** | **0.06** | **0.07** | 0.02 | **0.26** |
| NMB | 0.00 | 0.02 | 0.02 | 0.12 | 0.11 | 0.08 | 0.04 | 0.00 | **0.37** | **0.41** | **0.10** | **0.15** | **0.14** | **0.49** | **0.12** | **0.18** | 0.05 | **0.24** |
| NPB | 0.04 | 0.01 | 0.00 | 0.01 | 0.03 | 0.01 | 0.04 | 0.00 | **0.51** | **0.46** | **0.05** | **0.04** | **0.03** | **0.02** | **0.03** | **0.02** | 0.02 | **0.14** |
| LPL | 0.00 | 0.01 | 0.01 | 0.02 | 0.00 | 0.03 | 0.09 | 0.01 | **0.04** | **0.11** | **0.03** | **0.11** | **0.24** | **0.35** | **0.03** | **0.18** | 0.02 | **0.14** |
| PIF1 | 0.00 | 0.02 | 0.00 | 0.01 | 0.03 | 0.01 | 0.01 | 0.01 | **0.28** | **0.14** | **0.06** | **0.17** | **0.07** | **0.15** | **0.06** | **0.04** | 0.01 | **0.12** |
| GNG4 | 0.00 | 0.00 | 0.00 | 0.00 | 0.00 | 0.01 | 0.00 | 0.00 | **0.14** | **0.30** | **0.03** | **0.00** | **0.01** | **0.01** | **0.01** | **0.00** | 0.00 | **0.06** |
| KRT86 | 0.00 | 0.00 | 0.00 | 0.00 | 0.00 | 0.00 | 0.02 | 0.01 | **0.00** | **0.02** | **0.12** | **0.01** | **0.06** | **0.04** | **0.09** | **0.06** | 0.00 | **0.05** |
| ZBED2 | 0.00 | 0.00 | 0.00 | 0.00 | 0.00 | 0.00 | 0.01 | 0.00 | **0.01** | **0.03** | **0.01** | **0.02** | **0.07** | **0.01** | **0.00** | **0.00** | 0.00 | **0.02** |

Table S5. Pathological data of formalin-fixed, paraffin-embedded (FFPE) HBV-associated HCC samples.

| Hepatitis virus status | Gender | Age | Tumor nodules (#) | Tumor size (cm) | Tumor encapsulation | Venous invasion | Edmondson grade | Background liver | BCLC | HBV DNA (IU/mL) | IHC Positive Area (%) |
|---|---|---|---|---|---|---|---|---|---|---|---|
| HBV | male | 62 | 1 | 6 | complete | yes(small+ large vessels) | II | fibrosis | B | 279,811 | **89** |
| HBV | male | 58 | 1 | 3.3 | complete | no | II | chronic hepatitis | A | 99,668 | **40** |
| HBV | male | 52 | 1 | 3 | complete | no | II | chronic hepatitis | C | 5,343 | **55** |
| HBV | male | 46 | 3 | 1.3 | complete | no | II | chronic hepatitis | 0 | 616,000 | **25** |
| HBV | male | 75 | 1 | 12.7 | complete | no | II | chronic hepatitis | B | 152,000 | **20** |
| HBV | male | 65 | 1 | 1.1 | complete | no | II | chronic hepatitis | 0 | 127,000 | **20** |
| HBV | male | 50 | 1 | 4.7 | complete | no | II | fatty liver | A | 63,300 | **30** |
| HBV | male | 40 | 1 | 6 | complete | no | II | liver cirrhosis | A | 6,120,000 | **50** |
| HBV | male | 64 | 2 | 1.2+0.5 | complete | no | III | chronic hepatitis | C | 121,000 | **39** |
| HBV | male | 39 | 1 | 20 | complete | yes(small+ large vessels) | III | Chronic liver disease | C | 2,510,000 | **79** |
| HBV | male | 55 | 1 | 3 | complete | no | II | liver cirrhosis | A | 82,600 | **39** |
| HBV | male | 65 | 1 | 13.4 | | yes(small vessels) | II | chronic hepatitis | B | 44,400 | **25** |
| HBV | male | 49 | 1 | 20 | partial | yes(small+ large vessels) | III | chronic hepatitis | C | 2,247 | **90** |
| HBV | male | 48 | 1 | 1.5 | complete | no | II | liver cirrhosis | 0 | 5,048 | **9** |
| HBV | female | 64 | 1 | 10.5 | complete | yes(small vessels) | III | liver cirrhosis | C | 21,723 | **80** |
| HBV | male | 60 | 1 | 7.8 | partial | yes(small vessels) | II | liver cirrhosis | B | 1,573 | **60** |
| HBV | male | 63 | 2 | 3+1.6 | complete | no | II | liver cirrhosis | A | 2,944 | **0** |
| HBV | male | 62 | 1 | 5.5 | complete | yes(small vessels) | II | chronic hepatitis | B | 68,394 | **60** |
| HBV | male | 66 | 1 | 2.8 | complete | yes(small vessels) | II | chronic hepatitis | A | 8,116 | **90** |
| HBV | male | 71 | 3 | 1.6+3.2+2.5 | partial | no | II | liver cirrhosis | B | 242,974 | **90** |
| HBV | male | 58 | 1 | 11 | partial | yes(small vessels) | II | liver cirrhosis | B | 5,476,049 | **85** |
| HBV | male | 61 | 1 | 1.4+1.2 | partial | no | II | liver cirrhosis | A | 1,846,872 | **10** |
| NBNC | male | 60 | 1 | 3.5 | complete | yes(small+ large vessels) | II | normal liver | B | nil | **20** |
| NBNC | male | 64 | 2 | 6.5+2 | complete | yes(small vessels) | II | fatty liver | B | nil | **35** |
| NBNC | male | 72 | 1 | 17 | none | no | I | liver cirrhosis | B | nil | **20** |
| NBNC | male | 76 | 1 | 1.9 | complete | no | I | fibrosis | 0 | nil | **0** |
| NBNC | male | 76 | 1 | 6 | partial | no | II | liver cirrhosis | D | nil | **40** |
| NBNC | female | 37 | 1 | 2 | complete | no | II | normal liver | 0 | nil | **11** |
| NBNC | male | 69 | 1 | 6.1 | partial | no | II | normal liver | A | nil | **20** |
| NBNC | male | 64 | 1 | 10 | complete | no | II | normal liver | A | nil | **10** |
| NBNC | female | 57 | | 8.3 | none | yes(small vessels) | III | chronic hepatitis | C | nil | **0** |
| NBNC | male | 54 | 1 | 6.5 | complete | yes(small vessels) | II | fibrosis | B | nil | **40** |
| NBNC | female | 55 | 1 | 1.5 | partial | no | I | fatty liver | 0 | nil | **0** |
| NBNC | male | 46 | 2 | 2.7+1.1+1.5 | partial | no | II | liver cirrhosis | B | nil | **50** |
| NBNC | male | 60 | 1 | 1.9 | partial | no | II | liver cirrhosis | 0 | nil | **45** |
| NBNC | female | 62 | 1 | 3 | none | no | I | fatty liver | A | nil | **5** |
| NBNC | female | 66 | 1 | 2 | none | no | II | liver cirrhosis | 0 | nil | **9** |
| NBNC | female | 79 | 1 | 5 | complete | yes(small vessels) | II | normal liver | A | nil | **5** |
